# A Meta-Analysis of the Effects of Chronic Stress on the Prefrontal Transcriptome in Animal Models and Convergence with Existing Human Data

**DOI:** 10.1101/2025.10.23.683091

**Authors:** Jinglin Xiong, Megan H. Hagenauer, Cosette A. Rhoads, Elizabeth Flandreau, Nancy Rempel-Clower, Erin Hernandez, Duy Manh Nguyen, Annaka Saffron, Toni Duan, Stanley Watson, Huda Akil

## Abstract

**Background:** Chronic stress is a major risk factor for psychiatric disorders, including anxiety, depression, and post-traumatic stress disorder. Chronic stress can cause structural alterations like grey matter atrophy in key emotion-related areas such as the prefrontal cortex (PFC). To identify biological pathways affected by chronic stress in the PFC, researchers have performed transcriptional profiling (RNA-sequencing, microarray) to measure gene expression in rodent models. However, transcriptional signatures in the PFC that are shared across different chronic stress paradigms and laboratories remain relatively unexplored.

**Methods:** We performed a meta-analysis of publicly available transcriptional profiling datasets within the *Gemma* database. We identified six datasets that characterized the effects of either chronic social defeat stress (CSDS) or chronic unpredictable mild stress (CUMS) on gene expression in the PFC in mice (*n*=117). We fit a random effects meta-analysis model to the chronic stress effect sizes (log(2) fold changes) for each transcript (*n*=21,379) measured in most datasets. We then compared our results with two other published chronic stress meta-analyses, as well as transcriptional signatures associated with psychiatric disorders.

**Results:** We identified 133 genes that were consistently differentially expressed across chronic stress studies and paradigms (false discovery rate (FDR)<0.05). Fast Gene Set Enrichment Analysis (fGSEA) revealed 53 gene sets enriched with differential expression (FDR<0.05), dominated by glial and neurovascular markers (*e.g.,* oligodendrocyte, astrocyte, endothelial/vascular) and stress-related signatures (*e.g.,* major depressive disorder, hormonal responses). Immediate-early gene markers of neuronal activity (*Fos, Junb, Arc, Dusp1*) were consistently suppressed. Many of the identified effects resembled those seen in previous meta-analyses characterizing stress effects (CSDS, early life stress), despite minimal overlap in included samples. Moreover, some effects resembled previous observations from psychiatric disorders, including alcohol abuse disorder, major depressive disorder, bipolar disorder, and schizophrenia.

**Conclusion:** Our study demonstrates that chronic stress induces a robust, cross-paradigm PFC signature characterized by down-regulation of glia/myelin and vascular pathways and suppression of immediate-early gene activity, highlighting cellular processes linking chronic stress exposure, PFC dysfunction, and psychiatric disorders.

**Graphical Abstract:** 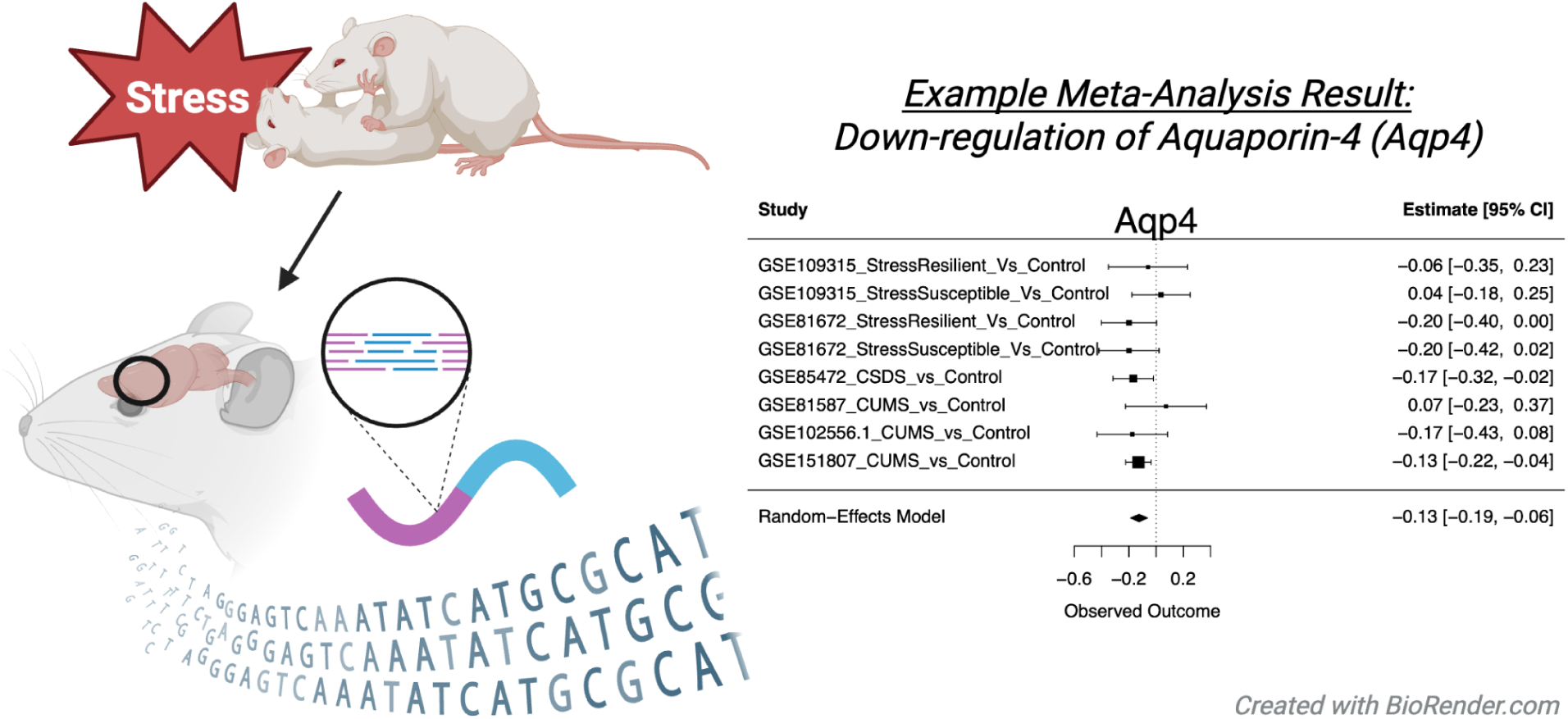

**Key Points:**

- Chronic stress is linked to many human psychiatric disorders involving the prefrontal cortex (PFC).
- Meta-analysis showed that stressed mice had less gene expression for glia and neural activity in the PFC.
- PFC gene expression in chronically stressed mice mirrors patterns seen in psychiatric patients.

**Plain Language Summary:** Chronic stress has been shown to have lasting effects on the brain, contributing to cognitive impairments and the risk of psychiatric disorders such as anxiety, depression, and post-traumatic stress disorder. The prefrontal cortex, a brain region that is important for behavioral and attentional control, is sensitive to chronic stress. We combined information from six public mouse datasets to identify consistent effects of chronic stress on gene expression (mRNA) in the prefrontal cortex. In these studies, mice that had experienced chronic stress showed a decreased amount of mRNA related to a variety of non-neuronal support cells (glia) and blood vessels, and general neural activity. The mouse chronic stress signature overlapped with gene-activity patterns reported in human psychiatric conditions, suggesting a biological bridge between chronic stress and the onset of psychopathology.

## Introduction

Stress is defined as the psychophysiological response triggered by an actual or perceived threat (Schneiderman et al., 2005). Acute stress usually does not pose a significant health risk - indeed, some activation of the stress system is essential at the neurobiological and functional level (McEwen & Akil, 2020). Chronic stress, however, can exceed the system’s capacity to return to homeostasis, resulting in high allostatic load coupled with structural alterations in key brain regions (Schneiderman et al., 2005; McEwen et al., 2016), cognitive impairment (Lupien et al., 2009), and the development of psychiatric disorders, including major depressive disorder (MDD) and post-traumatic stress disorder (PTSD) (Musazzi et al., 2023; Westfall et al., 2021).

To study chronic stress, researchers use diverse paradigms in laboratory rodent models including, but not limited to, social isolation, chronic mild stress, repeated restraint stress, chronic social defeat stress (CSDS), social instability stress, and chronic unpredictable/variable stress (CUMS)(Becker et al., 2021; Touchant & Labonté, 2022). Animals subjected to these different stress paradigms can manifest different responses but also show common behavioral phenotypes. For example, CSDS, social isolation, early life stress, and CUMS have all been shown to induce anxiety-like and depression-like behaviors (Nestler & Hyman, 2010; Touchant & Labonté, 2022), and both the CSDS and CUMS paradigms are widely used models for MDD (Gururajan et al., 2019; Nestler & Hyman, 2010). These models have revealed that susceptibility and resilience to stress-related emotional disorders are likely affected by an interaction of genetic inborn differences, epigenetic programs, and environmental factors (Clinton et al., 2022).

A key brain region in stress regulation is the prefrontal cortex (PFC). PFC is involved in emotional processing and cognition (Gray et al., 2002). Chronic stress causes structural and functional changes in the PFC, including retraction in the dendrites of pyramidal cells (Arnsten, 2009). Therefore, analyzing stress-induced changes in gene expression within the PFC may improve our understanding of the etiology of stress-sensitive disorders.

Transcriptional profiling, such as RNA sequencing (RNA-Seq) and microarray, can yield a large volume of information about the functional effects of stress-related interventions, yet also present many challenges, including large-scale technical noise (Hicks et al., 2018). These studies are often conducted with small sample sizes, which increases the likelihood of false negative and false positive results due to low statistical power (Button et al., 2013). To draw a more conclusive picture of stress-induced gene expression changes in the PFC, we conducted a meta-analysis of transcriptional profiling data from publicly available datasets investigating the effect of chronic stress in laboratory rodents.

To minimize bias, we used standardized and systematic methods to identify, extract, and exclude datasets and run the meta-analysis. These methods enabled us to identify PFC transcriptional signatures related to chronic stress that are consistent across experiments and chronic stress paradigms. To strengthen the validity of our findings, we compared our results with recent related meta-analyses and systematic reviews that employed different methodological approaches (Duan et al., 2025; Gururajan, 2022; Reshetnikov et al., 2022; Stankiewicz et al., 2022a). Additionally, we explored the relationship between chronic stress-induced transcriptional changes and those observed in psychiatric disorders by comparing our findings with large-scale meta-analyses of data from human cortical tissue (Gandal et al., 2018; Piras et al., 2022). These comparisons allowed us to examine the translational relevance of our findings and identify convergent stress-related molecular signatures.

## Methods

### Meta-Analysis Methods

#### General Overview

This meta-analysis was conducted as part of the *Brain Data Alchemy Project,* a collective effort to improve the reliability and generalizability of transcriptional profiling findings using meta-analyses of public datasets. This guided effort uses a standardized pipeline for dataset identification, inclusion/exclusion, and meta-analysis (protocol for 2022: (M. Hagenauer et al., 2024), example validation: (Rhoads et al., 2025), *not pre-registered*). It capitalizes on the data curation, preprocessing, and analysis efforts of the Gemma database (Lim et al., 2021; Zoubarev et al., 2012). Gemma currently houses >19,000 re-analyzed transcriptional profiling datasets, with preprocessing that includes up-to-date microarray probe alignment and RNA-Seq read mapping, sample and gene-level quality control, and identification of potential batch confounds. Within Gemma, differential expression analyses are performed using the *limma* (microarray) or *limma-voom* (RNA-Seq) pipeline, providing users with outputs from omnibus (full dataset) tests and individual contrasts. Our analysis code (R v.4.2.0, R studio v.2022.02.4) is released at: https://github.com/Jinglin0320/Metaanalysis_GemmaOutput_CSDS_PFC_Jinglin.

#### Dataset Identification and Selection

Potential datasets were identified in the Gemma database (https://gemma.msl.ubc.ca/home.html) using pre-specified search terms related to chronic stress paradigms (*n*=174 datasets), and narrowed using standardized inclusion/exclusion criteria (**Figure 1**). To keep the project manageable, we narrowed the datasets further to CSDS and CUMS paradigms because they are well-established and widely used to model MDD (Gururajan et al., 2019; Nestler & Hyman, 2010). Both paradigms have particularly strong construct validity for the chronic uncontrollable, unpredictable stress that increases risk for depression in humans, consistently resulting in depression-like symptoms that can be reversed by chronic administration of SSRIs (Deslauriers et al., 2018; Gyles et al., 2024). These two models have also been shown to exhibit overlapping effects on the transcriptome, proteome, and metabolome in other brain areas (Gui et al., 2021; Gyles et al., 2024). We excluded one study that used a novel CSDS paradigm tailored to females (Deonaraine et al., 2020), and one study that used a non-standard CSDS protocol with males, exposing a single intruder to three resident mice for 2 h/day for 6 days (Wang et al., 2019). Preprocessed gene expression data was confirmed to fall into a range typical for log(2) expression (RNA-seq: -5 to 12, microarray: 4 to 15). One dataset (GSE114224, (Chae et al., 2021)) was excluded because it had a highly restricted range (2.5-4), suggesting that the log-transformation was performed incorrectly when the dataset was imported into the Gemma database.

**Figure 1.**
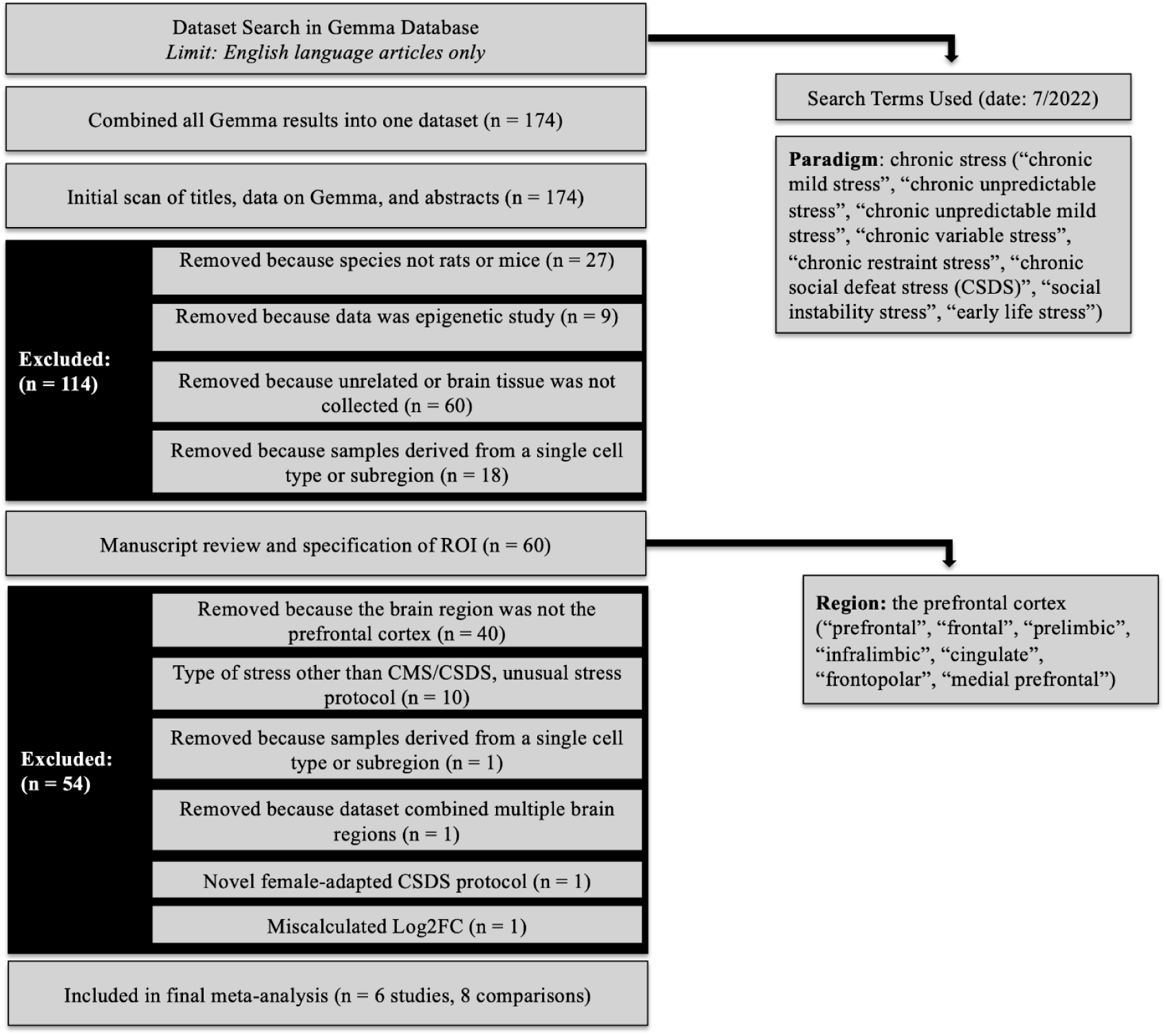
PRISMA diagram illustrating the dataset search terms and selection criteria used for the meta-analysis. Abbreviations: ROI=Region of Interest, Log2FC=Log(2)Fold Change, n= number of datasets.

#### Characteristics of the Selected Datasets

Six studies survived our criteria (*n*=8 stress vs. control contrasts, **Table 1**). Three studies utilized the CUMS paradigm, with animals exposed to alternating stressors for either three weeks (Ma et al., 2016; Labonté et al., 2017) or four weeks (Yu et al., 2021), one of which used behavioral testing to screen for stress susceptible animals (Ma et al., 2016). The other three studies employed the CSDS paradigm followed by a social interaction test to classify the mice into either a stress susceptible or stress resilient group. One of these studies administered CSDS for 14 days (Lehmann et al., 2017) and the other two studies applied CSDS for 10 days (Bagot et al., 2017; Laine et al., 2018). One study used 10-minutes of daily contact with the aggressor (Laine et al., 2018) and the other two used 5-minutes (Bagot et al., 2017; Lehmann et al., 2017). As our goal was to characterize generalized chronic stress responses, we included the effects of stress derived from both stress susceptible and stress resilient animals in our meta-analysis. However, due to common interest in the topic and potential clinical relevance (Nestler & Russo, 2024), we maintained separate effect size estimates for the susceptible and resilient groups, allowing readers to view any related heterogeneity in forest plots and clustering analyses. In general subjects were overwhelmingly male, with the one study that included both sexes (Labonté et al., 2017) coded with the reference level for sex set as male in the differential expression model in *Gemma*. Altogether, there was a final sample size of *n*=117 subjects, which should be sufficiently powered (80% power at alpha=0.05) to detect medium effect sizes.

**Table 1.**
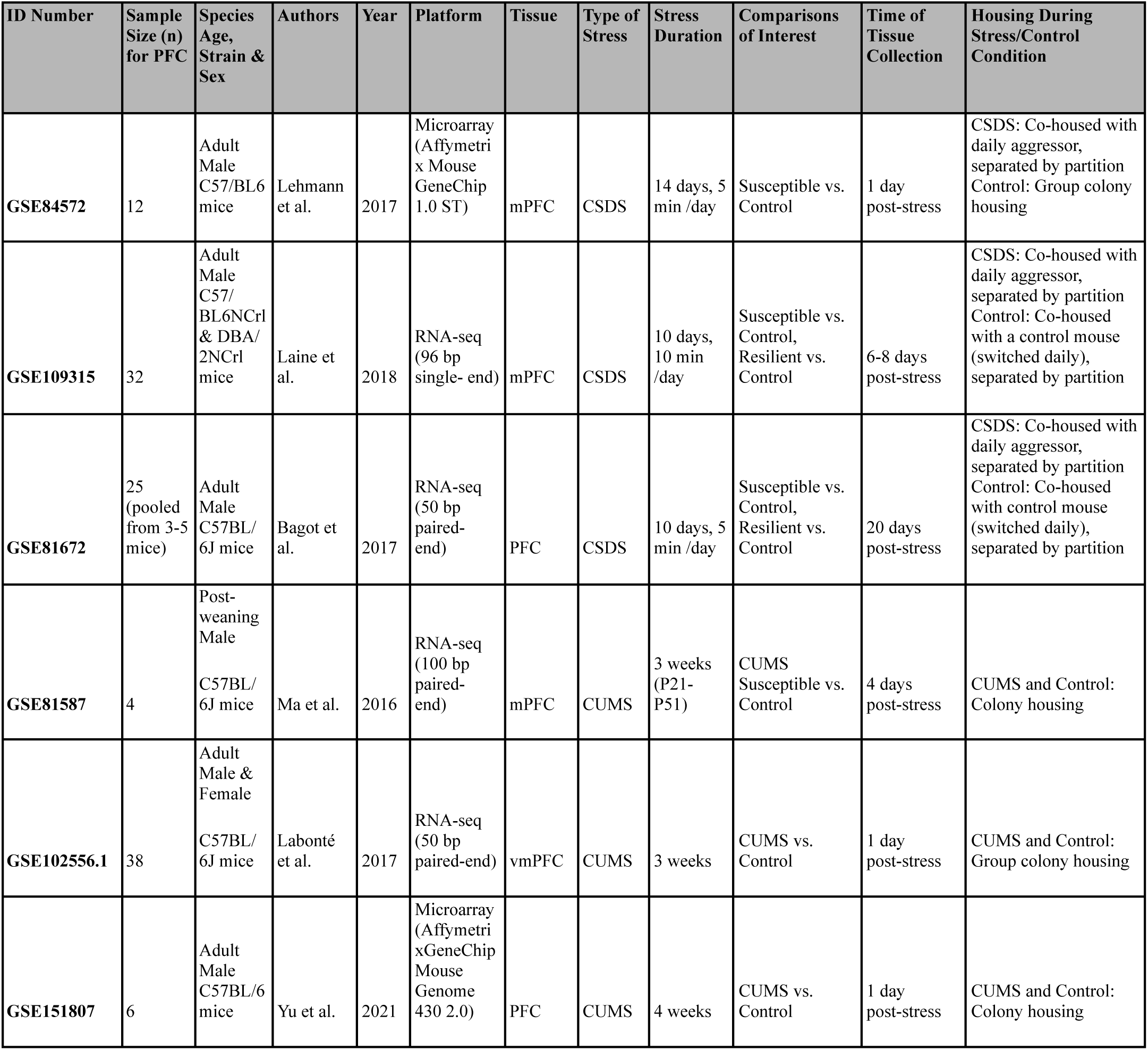
Overview of the datasets included in the meta-analysis. The table lists the Gene Expression Omnibus (GEO) ID for the dataset, sample size (n) for the PFC experiment, authors and year of publication (Bagot et al., 2016; Labonté et al., 2017; Laine et al., 2018; Lehmann et al., 2017; Ma et al., 2016; Yu et al., 2021), and transcriptional profiling platform. Study details include the species, strain, age, and sex of the subjects, brain region examined, the type of chronic stress, the duration of stress, specific experimental groups included in the statistical comparisons (contrasts) extracted for inclusion in the meta-analysis, delay between stress and tissue collection, and housing condition during the stress or control intervention. Abbreviations: PFC (prefrontal cortex, not otherwise defined), mPFC (medial PFC), vmPFC (ventromedial PFC), CSDS (chronic social defeat stress), CUMS (chronic unpredictable mild stress or chronic variable stress), Min: minute, P: postnatal age.

| ID Number | Sample Size (n) for PFC | Species Age, Strain & Sex | Authors | Year | Platform | Tissue | Type of Stress | Stress Duration | Comparisons of Interest | Time of Tissue Collection | Housing During Stress/Control Condition |
| --- | --- | --- | --- | --- | --- | --- | --- | --- | --- | --- | --- |
| GSE84572 | 12 | Adult Male C57/BL6 mice | Lehmann et al. | 2017 | Microarray (Affymetri x Mouse GeneChip 1.0 ST) | mPFC | CSDS | 14 days, 5 min /day | Susceptible vs. Control | 1 day post-stress | CSDS: Co-housed with daily aggressor, separated by partition<br>Control: Group colony housing |
| GSE109315 | 32 | Adult Male C57/BL6NcrI & DBA/2NcrI mice | Laine et al. | 2018 | RNA-seq (96 bp single- end) | mPFC | CSDS | 10 days, 10 min /day | Susceptible vs. Control, Resilient vs. Control | 6-8 days post-stress | CSDS: Co-housed with daily aggressor, separated by partition<br>Control: Co-housed with a control mouse (switched daily), separated by partition |
| GSE81672 | 25 (pooled from 3-5 mice) | Adult Male C57BL/6J mice | Bagot et al. | 2017 | RNA-seq (50 bp paired-end) | PFC | CSDS | 10 days, 5 min /day | Susceptible vs. Control, Resilient vs. Control | 20 days post-stress | CSDS: Co-housed with daily aggressor, separated by partition<br>Control: Co-housed with control mouse (switched daily), separated by partition |
| GSE81587 | 4 | Post-weaning Male C57BL/6J mice | Ma et al. | 2016 | RNA-seq (100 bp paired-end) | mPFC | CUMS | 3 weeks (P21-P51) | CUMS Susceptible vs. Control | 4 days post-stress | CUMS and Control: Colony housing |
| GSE102556.1 | 38 | Adult Male & Female C57BL/6J mice | Labonté et al. | 2017 | RNA-seq (50 bp paired-end) | vmPFC | CUMS | 3 weeks | CUMS vs. Control | 1 day post-stress | CUMS and Control: Group colony housing |
| GSE151807 | 6 | Adult Male C57BL/6 mice | Yu et al. | 2021 | Microarray (Affymetri xGeneChip Mouse Genome 430 2.0) | PFC | CUMS | 4 weeks | CUMS vs. Control | 1 day post-stress | CUMS and Control: Colony housing |

#### Result Extraction

The differential expression results for each of the relevant eight stress-related statistical contrasts (stress vs. control) were imported into R. Rows with probes or transcript sequences that mapped to multiple gene annotations (official gene symbol) were excluded, and gene-level average effect sizes (log(2) fold change or Log2FC), standard errors (SE), and sampling variances (SV) were calculated and aligned across datasets by gene symbol.

#### Meta-Analysis

To perform the meta-analysis, we used the *metafor* package (v.3.4.0, (Viechtbauer, 2010)) to fit a random effects model to the effect sizes of chronic stress (Log2FC values) and accompanying SV for each gene represented in at least five stress vs. control statistical contrasts (*n*=21,379 genes). A minimum of five stress vs. control contrasts was chosen to ensure that a minimum of at least three studies were represented in the meta-analysis, to increase the power and generalizability of the results. Random effects modelling was chosen to account for the heterogeneous methods and sample characteristics across the included studies that might introduce variability amongst their true effects. To fit the random effects model, we used the function *rma()* from the *metafor* package, which treats this heterogeneity as purely random (normally distributed), estimated using restricted maximum-likelihood estimation (REML). The function uses the inverse-variance method, which treats the precision of each study’s estimated effect as inversely related to the study’s SV. Due to limited dataset availability, we used a simple, intercept-only model as our main outcome. The resulting p-values were corrected for false discovery rate (FDR) using the Benjamini-Hochberg method (*multtest* package: v.2.8.0, (Pollard et al., 2005)).

#### Functional Ontology

To identify functional patterns, fast Gene Set Enrichment Analysis (*fGSEA:* (Sergushichev, 2016)) was performed on the meta-analysis results, ranked by estimated Log2FC, using Brain.GMT (M. H. Hagenauer et al., 2024). Brain.GMT is a curated database of gene sets related to nervous system function, tissue, and cell types, packaged with traditional gene ontology gene sets (MSigDB “C5”: 14,996 gene sets).

### Comparison of Meta-Analysis Results to Related Studies

#### Other Meta-Analyses Examining Chronic Stress Effects on the Rodent PFC

While conducting our study, several meta-analyses were published addressing the effects of chronic stress on the PFC. These included a meta-analysis of the effects of either 10 days or 30 days of CSDS (Reshetnikov et al., 2022), a small meta-analysis of stress-susceptibility vs. stress resilience (Reshetnikov et al., 2022), and a meta-analysis of the effects of chronic stress, more broadly defined (CSDS, CUMS, or chronic restraint) (Gururajan, 2022). Although these meta-analyses used public data, due to focusing exclusively on RNA-Seq datasets and differing inclusion/exclusion criteria they showed either only small partial overlap with our own sample (*10 days CSDS*: n=56, overlap 15%: n=18 of our 117; *stress susceptibility vs. resilience:* overlap 15%: n=18 of our 117; *Chronic Stress:* n=101, overlap 32%: 37 of our 117) or no overlap (*30 days CSDS:* n=20), and therefore could provide some independent insight into our findings. Results from the CSDS meta-analyses were extracted from *Table STS3A* (*10 Days CSDS*), *Table STS3C* (stress susceptibility vs. resilience), and *Table STS3B* (*30 Days CSDS*) (Reshetnikov et al., 2022), which only contained results for genes surviving a nominal *p*<0.05 threshold. Results from the chronic stress meta-analysis were extracted from *Suppl Table 1G* (*“Combined Analysis”*) (Gururajan, 2022), which only contained results for genes surviving an FDR<0.05 threshold.

#### Other Large Studies Characterizing Stress Effects on the Rodent Brain

We also compared our results to other studies examining stress effects on the brain using large sample sizes. These included a meta-analysis of the effects of early life stress on the PFC transcriptome ((Duan et al., 2025): *n*=89, full results for all genes extracted from *Table S1*) and a large study examining the effects of chronic daily corticosterone on the hippocampal transcriptome ((Juszczak et al., 2025): *n*=48, full results for all genes extracted from *Suppl Data 2 (“All”)).* The supplement for (Juszczak et al., 2025) also provided a comprehensive database including results from large*“vote-counting”-*style literature reviews characterizing the direction of effect (up, down, mixed effects) for genes showing reported effects of acute and chronic stress in brain transcriptomic studies (Stankiewicz et al., 2022): 79 studies reviewed).

We also ran several exploratory analyses that are more tangential and relegated to the supplement: We ran comparisons with the effects of acute stress and corticosterone treatment (Jaszczyk et al., 2023; Juszczak et al., 2025; Stankiewicz et al., 2022a) and acute sleep deprivation (Diessler et al., 2018; Rhoads et al., 2025). We also conducted a small follow-up analysis to explore whether our meta-analysis findings might be male-specific using comparisons to published differential expression results broken down by sex (Deonaraine et al., 2020; Labonté et al., 2017; Shao et al., 2025).

#### Large Studies Characterizing the Effects of Psychiatric Disorders on the Cortex

We compared our results to the effects of multiple psychiatric disorders on the cortical transcriptome. These included several large meta-analyses examining the effects of suicide in different regions of the PFC (*orbitofrontal PFC, dorsolateral PFC, PFC region unspecified*, (Piras et al., 2022): *n*=380). Results were extracted from *Table S1*, which contained results for genes surviving an FDR<0.05 threshold in any of the four meta-analyses. We also referenced large microarray meta-analyses characterizing the effects of multiple psychiatric disorders on the cortex (MDD (BA9, BA25, BA46: *n*=176), Bipolar Disorder (BA46, parietal: *n=*197), and Schizophrenia (BA46, BA10, parietal: *n*=314)), and a re-analysis of the effects of Alcohol Abuse Disorder on the superior frontal cortex (n=32) (*Table S1* in (Gandal et al., 2018): results for all genes), as well as two large validation meta-analyses of RNA-Seq data characterizing the effects of Bipolar Disorder (*n*=171) and Schizophrenia (*n*=384) in the frontal cortex (*Table S2* in (Gandal et al., 2018): results for all genes). It is important to note that the results of these psychiatric meta-analyses can only be considered partially independent: the Bipolar Disorder and Schizophrenia microarray meta-analyses share an overlapping set of control subjects, which also partially overlap the MDD microarray meta-analysis. Likewise, the Bipolar Disorder and Schizophrenia RNA-Seq meta-analyses share an overlapping set of control subjects. In contrast, there was no overlap between the samples included in the large meta-analyses of (Gandal et al., 2018) and (Piras et al., 2022). We also compared our meta-analysis results to a review of reported PTSD effects on the PFC transcriptome ((Stankiewicz et al., 2022): *n*=121), summarized in terms of categorical direction of effect (*Suppl Data 2* of (Juszczak et al., 2025)).

To compare our results with human findings, we extracted information about human and mouse gene orthology from “HCOP: Orthology Predictions Search” (https://www.genenames.org/tools/hcop/, bulk download: 8-21-2025). This information was filtered down to 1) one-to-one mouse-to-human mapped orthologs, 2) orthologs with matching gene symbols, and 3) orthologs with multiple-to-one mouse-to-human mappings that had the most databases supporting the orthology (*column: “support”*, ties broken by row index).

#### Statistical Methods for Result Comparisons

To conduct these comparisons, we extracted all available results for our 133 differentially expressed genes (“DEGs” FDR<0.05). If there were Log2FC values available, we examined the correlation with the Log2FC estimated from our chronic stress meta-analysis using both parametric (regression, Pearson’s correlation) and non-parametric (Spearman’s correlation) methods (functions: *summary.lm, cor.test*).

## Results

### Meta-Analysis

#### Overview

Of the 21,379 genes included in the meta-analysis, 21,296 produced stable meta-analysis estimates (*full results:* **Table S1**). Our meta-analysis revealed 133 DEGs (FDR<0.05), 97 of which were downregulated and 36 upregulated. A hierarchically-clustered heatmap of the top DEGs did not reveal a clear pattern related to stress paradigm or stress susceptibility, but potentially some clustering by platform (**Figure 2**).

**Figure 2.**
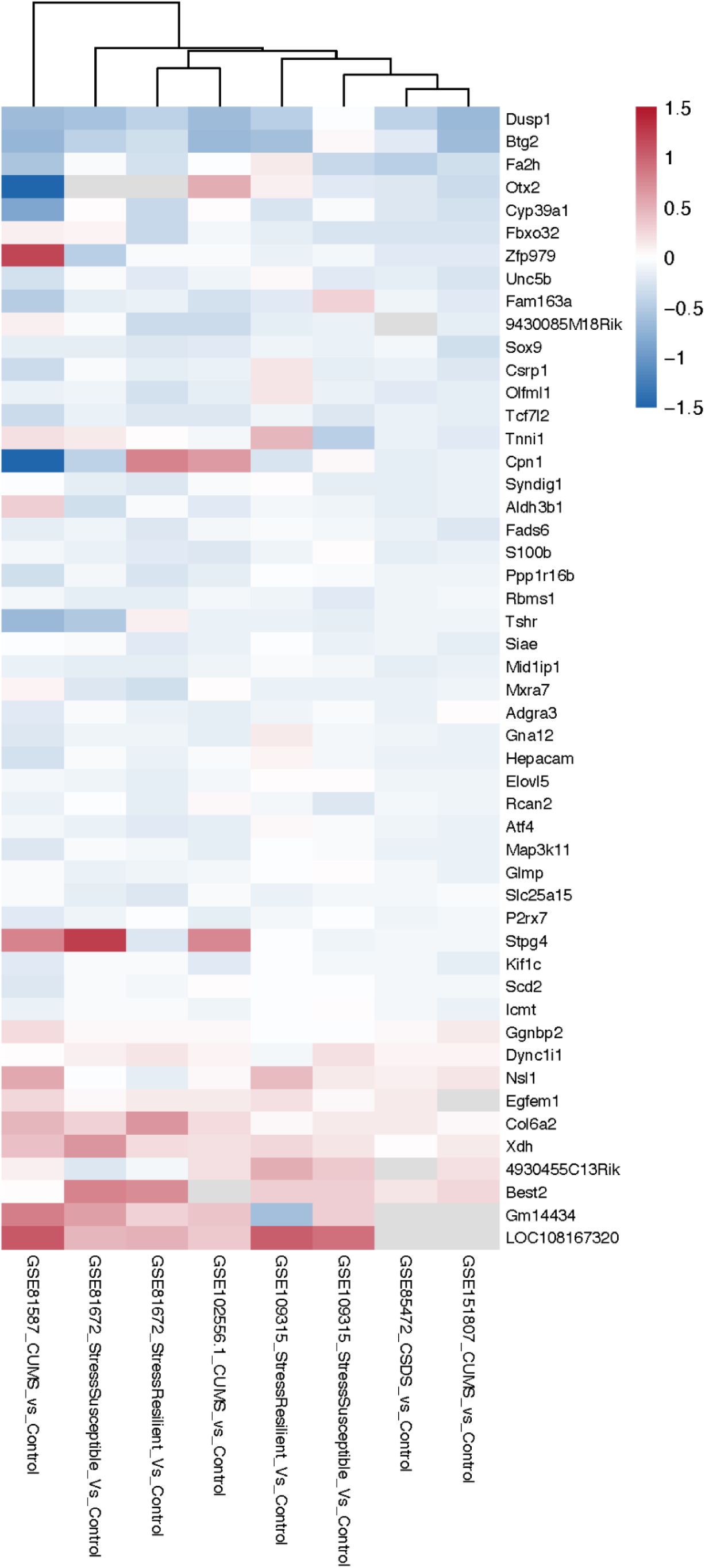
Heatmap of the top 50 differentially expressed genes across all 8 chronic stress vs. control comparisons. Rows correspond to the 50 genes with the smallest FDR, columns indicate the Gene Expression Omnibus (GEO) dataset ID (GSE#…) and specific chronic stress vs. control comparison. Columns are ordered by unsupervised hierarchical clustering of log2 fold-change (Log2FC) values so that chronic stress vs. control comparisons (contrasts) with similar results appear adjacent. The dendrogram at the top of the figure represents that similarity: the most similar items are joined at lower levels (“branches”), and less similar groups are joined higher up on the tree. Within the heatmap, pink indicates upregulation (positive Log2FC), and blue indicates down-regulation (negative Log2FC). Grey indicates that the gene lacks representation in the study.

Example DEGs are illustrated with forest plots, showing the consistency of differential expression across studies and paradigms (**Figures 3&4).** Chronic stress broadly downregulated gene expression important for glial development and function in the PFC (**Figure 3**), including Aquaporin 4 (*Aqp4*), S100 calcium-binding protein B (*S100b*), Fatty Acid 2-Hydroxylase (*Fa2h*), P2X purinoceptor 7 (*P2rx7*), ELOVL fatty acid elongase 5 (*Elovl5*), and Hepatocyte cell adhesion molecule (*Hepacam*). Chronic stress also suppressed immediate early gene (IEG) responses linked to neuronal activity and plasticity in the PFC (**Figure 4**), including Fos proto-oncogene, AP-1 transcription factor subunit (*Fos*), JunB proto-oncogene, AP-1 transcription factor subunit (*Junb*), Activity-regulated cytoskeleton-associated protein (*Arc*), and Dual specificity phosphatase 1 (*Dusp1*).

**Figure 3.**
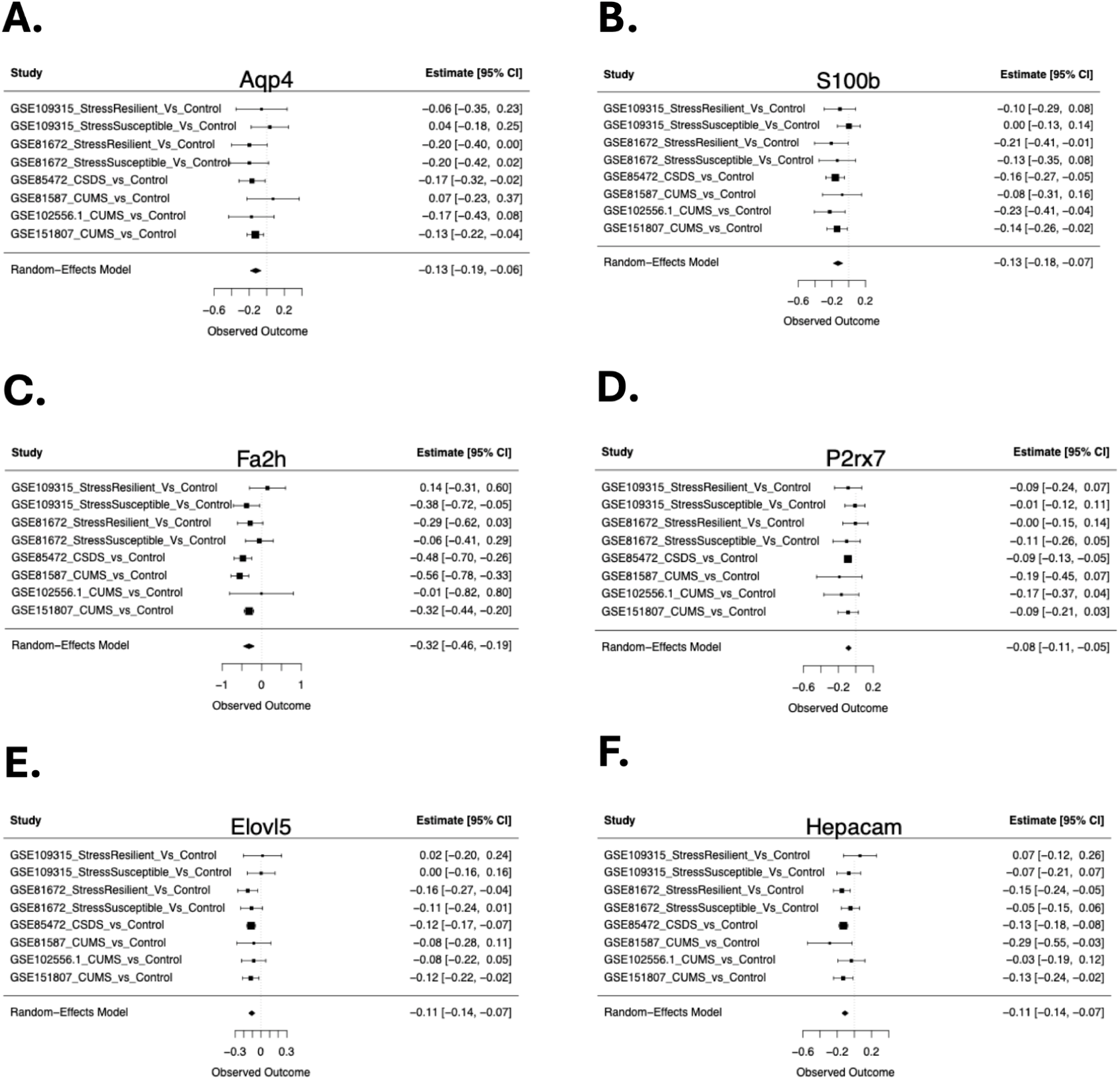
Genes related to glial development and function are down-regulated in the PFC in chronic stress models. Forest plots illustrate the consistency of the down-regulation across datasets and conditions. Rows illustrate Log2FC (squares) with 95% confidence intervals (whiskers) for each of the datasets and the meta-analysis random effects model (RE Model). **(A)** Forest plot for Aquaporin 4 (Aqp4). **(B)** Forest plot for S100 calcium-binding protein B (S100b). **(C)** Forest plot for Fatty acid 2-hydroxylase (Fa2h). **(D)** Forest plot for P2X purinoceptor 7 (P2rx7). **(E)** Forest plot for ELOVL fatty acid elongase 5 (Elovl5). **(F)** Forest plot for hepatocyte cell adhesion molecule (Hepacam).

**Figure 4.**
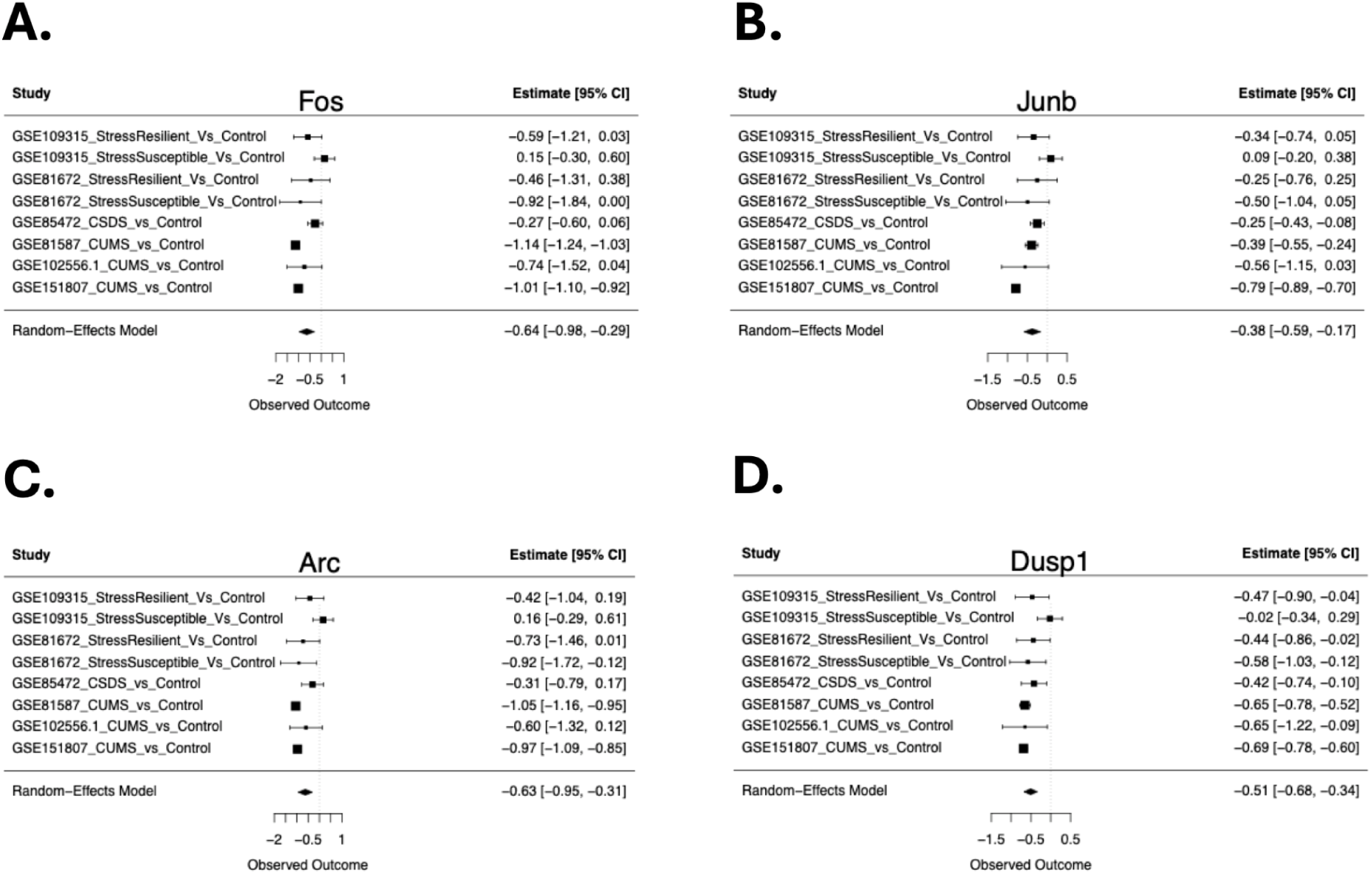
Immediate early genes (IEGs) are down-regulated in the PFC in chronic stress models. Forest plots illustrate the consistency of the down-regulation across datasets and conditions. Formatting follows the conventions of Figure 3. **(A)** Forest plot for Fos proto-oncogene, AP-1 transcription factor subunit (Fos). **(B)** Forest plot for JunB proto-oncogene, AP-1 transcription factor subunit (Junb). **(C)** Forest plot for Activity-regulated cytoskeleton-associated protein (Arc). **(D)** Forest plot for Dual specificity phosphatase 1 (Dusp1).

#### Functional Patterns

We performed a fast gene set enrichment analysis (*fGSEA*) using a database containing both brain-related (*Brain.GMT*) and traditional gene ontology gene sets. Of the 10,847 gene sets included in the results, 49 were enriched with down-regulation following chronic stress and 4 enriched with upregulation (FDR<0.05, **Table S2**).

Chronic stress altered gene sets that are associated with several brain cell types (**Table 2**), with glial cells appearing to be central responders. The most prominent changes involved oligodendrocytes (8 out of 53) and developing oligodendrocytes (4 out of 53), which were uniformly downregulated. Gene sets related to astrocytes and astrocytic functions (4 out of 53), radial-glia-like signatures (3 out of 53), and vasculature (4 out of 53) were also downregulated after chronic stress. Only one neuronal-related gene set was affected, and one gene set broadly related to cell differentiation was downregulated.

**Table 2.**
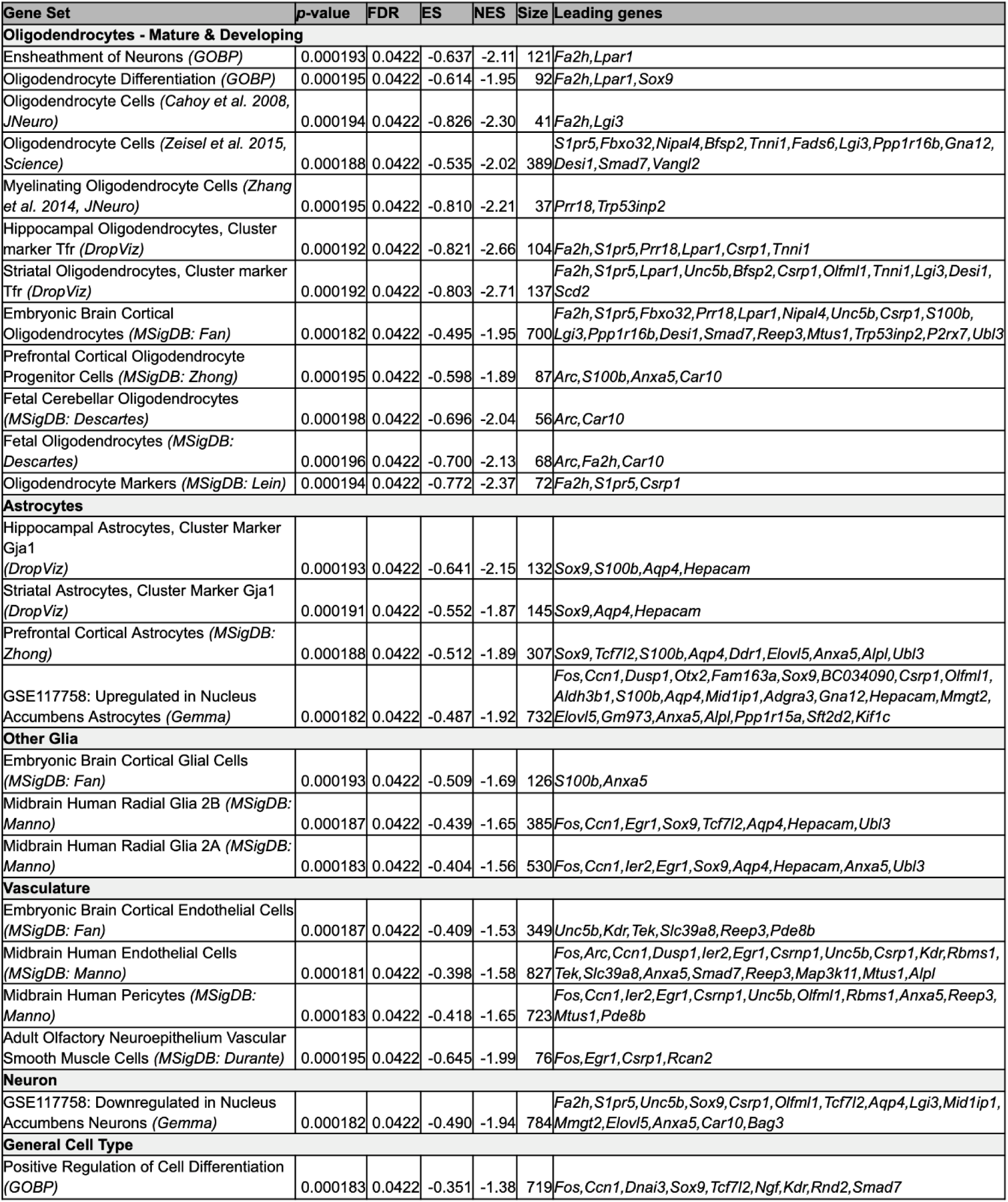
Functional gene sets that are enriched with the effects of chronic stress are related to various cell types. Size: Size of gene set. Leading genes: A list of the leading genes in the gene set that were significantly differentially expressed following chronic stress (FDR < 0.05) in the meta-analysis. Abbreviations: ES, enrichment score; FDR, false discovery rate; NES, normalized enrichment score. Gene set names have been reformatted for clarity.

Our meta-analysis results showed an enrichment of differential expression in gene sets representing a variety of stress paradigms (**Table 3**), providing some cross-validation. Notably, we observed enrichment in gene sets derived from CSDS experiments that were not included in our meta-analysis (*GSE72343,* PFC-specific results unavailable within the Gemma database (Bagot et al., 2016); *GSE149195:* PFC not profiled (DiSabato et al., 2021)). This convergence suggests that the molecular program we identified captures core elements of the brain’s chronic stress response that can be generalized to other brain regions. There was also enrichment in gene sets from other stress paradigms, including contextual fear conditioning, trace fear conditioning, forced swim stress, restraint stress, and novel stress. Some gene sets showed an enrichment of differential expression in our meta-analysis that showed the same direction of effect as the original experiment, whereas other stress gene sets were enriched but shifted in the opposite direction. These oppositely shifting gene sets were largely driven by classic IEGs including *Fos*, *Arc*, *Dusp1*, *Egr1.* We also observed an enrichment in gene sets related to hormonal response and MDD, highlighting the relevance of our findings to stress-related psychopathology.

**Table 3.** Functional gene sets that are enriched with the effects of chronic stress are related to various types of stress, hormonal response, and MDD. Size: Size of gene set. Leading genes: A list of the leading genes in the gene set that were significantly differentially expressed following chronic stress (FDR<0.05) in the meta-analysis. Abbreviations: ES, enrichment score; FDR, false discovery rate; NES, normalized enrichment score. Gene set names have been reformatted for clarity.

| Gene Set | p-value | FDR | ES | NES | Size | Leading genes |
| --- | --- | --- | --- | --- | --- | --- |
| **Stress: Contradictory Direction of Effect** |  |  |  |  |  |  |
| GSE100236: Genes upregulated by swim stress in the hippocampus (*Gemma*) | 0.000192 | 0.0422 | -0.716 | -2.30 | 99 | *Fos, Arc, Ccn1, Dusp1, Btg2, Ier2, Egr1, Csrnp1, Ppp1r15a* |
| GSE100236: Genes upregulated by novelty stress in the hippocampus (*Gemma*) | 0.000193 | 0.0422 | -0.666 | -2.19 | 115 | *Fos, Arc, Ccn1, Dusp1, Btg2, Ier2, Egr1, Csrnp1, Ppp1r15a* |
| GSE100236: Genes upregulated by restraint stress in the hippocampus (*Gemma*) | 0.000192 | 0.0422 | -0.682 | -2.23 | 112 | *Fos, Arc, Ccn1, Dusp1, Btg2, Ier2, Egr1, Csrnp1, Ppp1r15a* |
| GSE16158: Genes upregulated by trace fear conditioning in the hippocampus, timepoint 0.5h (*Gemma*) | 0.000195 | 0.0422 | -0.901 | -2.16 | 20 | *Fos, Arc, Dusp1, Btg2, Egr1* |
| GSE16158: Genes upregulated by trace fear conditioning in the hippocampus, timepoint 2h (*Gemma*) | 0.000195 | 0.0422 | -0.876 | -2.10 | 20 | *Fos, Arc, Dusp1, Egr1* |
| GSE50423: Genes upregulated by contextual fear conditioning in the hippocampus (*Gemma*) | 0.000193 | 0.0422 | -0.929 | -1.93 | 10 | *Fos, Arc, Dusp1, Btg2* |
| GSE63412: Genes upregulated by contextual fear conditioning memory retrieval (*Gemma*) | 0.000193 | 0.0422 | -0.932 | -2.04 | 13 | *Fos, Arc, Ccn1, Dusp1, Btg2* |
| GSE63412: Genes upregulated by contextual fear conditioning (*Gemma*) | 0.000193 | 0.0422 | -0.932 | -2.04 | 13 | *Fos, Arc, Ccn1, Dusp1, Btg2* |
| GSE74966: Genes upregulated by contextual fear conditioning (*Gemma*) | 0.000194 | 0.0422 | -0.659 | -2.09 | 90 | *Fos, Egr1, Kdr* |
| GSE74966: Genes upregulated by contextual fear conditioning and electric shock (*Gemma*) | 0.000195 | 0.0422 | -0.662 | -2.10 | 92 | *Fos, Egr1, Kdr* |
| **Stress: Confirmatory Direction of Effect** |  |  |  |  |  |  |
| GSE50423: Genes downregulated by contextual fear conditioning in the hippocampus, timepoint 4h (*Gemma*) | 0.000184 | 0.0422 | -0.423 | -1.61 | 464 | *Fos, Arc, Ccn1, Dusp1, Btg2, Egr1, Atf4* |
| GSE50423: Genes downregulated by contextual fear conditioning in the hippocampus, timepoint 24h | 0.000191 | 0.0422 | -0.517 | -1.79 | 173 | *Fos, Arc, Ccn1, Dusp1, Btg2, Egr1* |
| GSE72343: Genes upregulated in stress resilient mice following chronic social defeat stress (*Gemma*) | 0.000205 | 0.0422 | 0.779 | 2.21 | 44 | *Xdh* |
| GSE72343: Genes upregulated in stress susceptible mice following chronic social defeat stress (*Gemma*) | 0.000205 | 0.0422 | 0.779 | 2.21 | 44 | *Xdh* |
| GSE149195: Genes upregulated following social defeat stress (*Gemma*) | 0.000206 | 0.0422 | 0.791 | 2.02 | 26 |  |
| **Stress-Related:** |  |  |  |  |  |  |
| Response to Hormone (*GOBP*) | 0.000183 | 0.0422 | -0.348 | -1.38 | 763 | *Fos, Dusp1, Btg2, Egr1, Fbxo32, Cpn1, Tshr, Tek, Alpl* |
| Genes downregulated in Major Depressive Disorder (*MSigDB: Aston*) | 0.000191 | 0.0422 | -0.570 | -1.94 | 148 | *Lpar1, P2rx7* |
| GSE95740: Genes upregulated in the dentate gyrus following environmental enrichment (*Gemma*) | 0.000195 | 0.0422 | -0.608 | -1.87 | 74 | *Fos, Arc, Dusp1, Csrnp1, Syndig1* |

### Similarities with Previous Studies Examining the Effects of Stress & Glucocorticoids

Some of our DEGs showed effects resembling those reported in previous meta-analyses examining the effects of chronic stress on the PFC transcriptome. For example, a meta-analysis of the effects of 10 days of CSDS ((Reshetnikov et al., 2022): *n*=56) provided results for 1968 genes surviving a nominal *p*<0.05 threshold, including 21 of our 133 DEGs. For these 21 genes, there was a strong positive correlation with the Log2FCs from our meta-analysis (*R*=0.73, *p*=0.000147; *Rho*=0.78, *p*=4.32e-05, **Figure 5A**), despite minimal overlap in included samples (15%: *n*=18 of our 117 subjects). Whether this strong positive correlation would persist for our DEGs with unavailable results is unknown. This study also included a smaller meta-analysis comparing the subset of animals identified as stress susceptible vs. stress resilient ((Reshetnikov et al., 2022): *n*=18 subjects) which provided results for 2614 genes that survived a nominal p<0.05 threshold, including 30 of our 133 DEGs. For these 30 genes, there was also a strong positive correlation with the Log2FCs from our chronic stress meta-analysis (R=0.74, p=3.05e-06, Rho=0.77, p=2.16e-06, **Figure 5C**), providing preliminary evidence suggesting that susceptible animals may have generalized stress effects in the PFC that are amplified compared to resilient animals.

**Figure 5.**
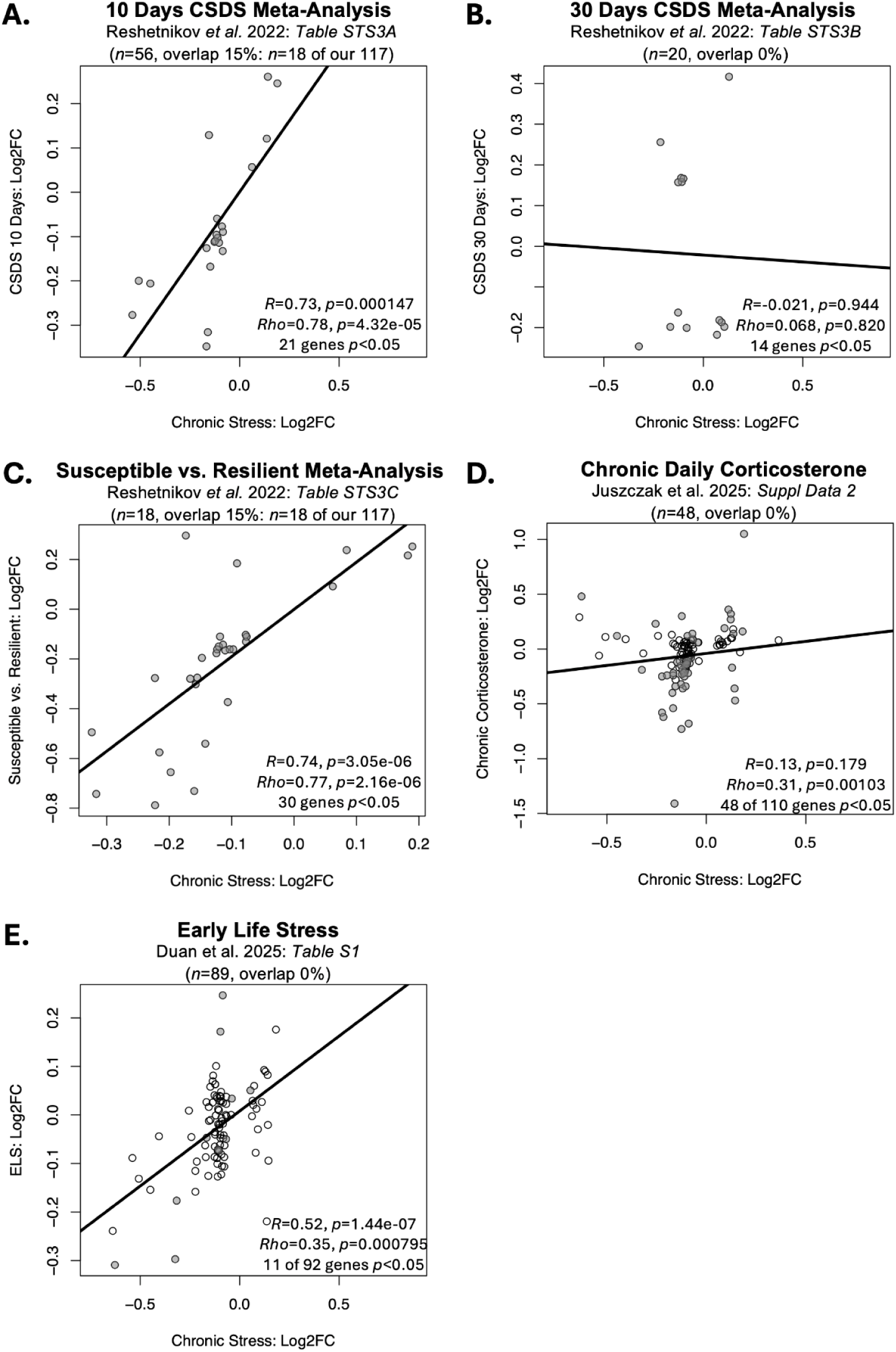
The DEGs from our current meta-analysis show similar effects in previous large studies characterizing the effects of chronic stress and glucocorticoids on the brain transcriptome. A-D: Scatterplots illustrate the correlation between the effects of chronic stress within our meta-analysis (Log2FC: x-axis) and the effects of stress within previous studies (Log2FC: y-axis) for any of our 133 identified DEGs (FDR<0.05) that had results available in the referenced publication. Data points are shaded grey if the effects within the previous publication were nominally significant (p<0.05). Trend lines illustrate the relationship between the two sets of results as identified by simple linear regression. **A.** Differential expression results were available for 21 of our DEGs within a previously published meta-analysis of the effects of 10 days of CSDS on the PFC ((Reshetnikov et al., 2022): n=56 subjects, only nominally significant results available). For these genes, there was a strong positive correlation with the Log2FCs in our meta-analysis (R=0.73, Rho=0.78). **B.** Differential expression results were available for 14 of our DEGs within a small previously published meta-analysis of the effects of 30 days of CSDS on the PFC ((Reshetnikov et al., 2022): n=20 subjects, only nominally significant results available). For these genes, there was minimal correlation between the Log2FCs from the two meta-analyses (R=-0.021, Rho=0.068). **C.** Differential expression results were available for 30 of our DEGs within a small previously published meta-analysis comparing animals identified as stress susceptible vs. stress resilient ((Reshetnikov et al., 2022): n=18 subjects, only nominally significant results available). For these genes, there was a strong positive correlation with the Log2FCs from our chronic stress meta-analysis (R=0.74, Rho=0.77). **D.** Differential expression results were available for 110 of our DEGs within a large study of the effects of chronic daily corticosterone on the hippocampal transcriptome ((Juszczak et al., 2025): n=48 subjects). For these genes, there was a small positive correlation with the Log2FCs in our meta-analysis (R=0.13, Rho=0.31). **E.** Differential expression results were available for 92 of our DEGs within a meta-analysis of the long-term effects of early life stress on the prefrontal transcriptome ((Duan et al., 2025): n=89 subjects). For these genes, there was a positive correlation with the Log2FCs in our meta-analysis (R=0.52, Rho=0.35).

Similar parallels were not observed with the results from a small meta-analysis of the effects of 30 days of CSDS ((Reshetnikov et al., 2022), *n*=20). Only 14 DEGs from our meta-analysis were represented in their provided results (3132 genes surviving a *p*<0.05 threshold), and their associated Log2FCs showed minimal correlation with our findings (*R*=-0.021, *p*=0.944; *Rho*=0.068, *p*=0.820, **Figure 5B**). We also observed limited overlap with the results from a meta-analysis of the effects of chronic stress more broadly defined (CSDS, CUMS, or chronic restraint) ((Gururajan, 2022), *n*=101). For this study, only results for genes surviving a stricter FDR<0.05 threshold were available (160 genes), of which only one (*Xdh*) was included in our DEGs, making comparison difficult.

We were able to more unambiguously document a positive correlation with results from large studies related to chronic stress with result reporting for more genes. For example, results were available for 92 of our DEGs within a meta-analysis of the long-term effects of early life stress on the prefrontal transcriptome ((Duan et al., 2025): *n*=89), eleven of which showed nominal significance (*p*<0.05). The Log2FCs for these genes correlated positively with the Log2FCs in our meta-analysis (92 genes: *R*=0.52, *p*=1.44e-07; *Rho*=0.35, *p*=0.000795, **Figure 5E**). Our results were also weakly correlated with the effects of chronic daily corticosterone on the hippocampal transcriptome ((Juszczak et al., 2025): *n*=48, **Figure 5D**), with 110 of our genes represented in their results, 48 of which showed nominal significance (*p*<0.05). For these genes, the Log2FCs showed a small positive correlation with the Log2FCs in our meta-analysis (110 genes: *R*=0.13, *p*=0.179; *Rho*=0.31, *p*=0.00103). Additional DEGs had previous relationships with chronic stress (either medium duration or prolonged stress) documented in a large review of the literature characterizing the effects of stress on the brain transcriptome (Stankiewicz et al., 2022) (**Table S3**). Altogether, these comparisons suggest that our findings add to a growing body of results documenting the effects of stress on brain gene expression.

### Similarities with Previous Studies Examining the Effects of Stress-Related Psychiatric Disorders

Chronic stress contributes to the onset and the exacerbation of many psychiatric disorders (Davis et al., 2017) and can induce anxiety-like and depression-like behaviors in animals (Deslauriers et al., 2018; Gyles et al., 2024). Chronic stress is also a consistent feature of life with a serious psychiatric disorder. Therefore, we decided to compare our meta-analysis results to previous findings from subjects with a variety of psychiatric disorders. We found that some of our DEGs showed effects resembling those reported in previous meta-analyses examining the effects of stress-related psychiatric disorders on the PFC transcriptome. Of 221 genes previously found to be differentially expressed in meta-analyses of the effects of suicide in the PFC (Piras et al., 2022), seven were amongst our 133 DEGs, all of which showed the same direction of effect. Amongst 40 genes that had previously shown consistent differential expression in association with PTSD in the PFC (Stankiewicz et al., 2022), three were amongst our DEGs, all of which also showed the same direction of effect.

When examining the Log2FCs for our DEGs, we also found a consistent positive correlation with the Log2FCs identified in previous meta-analyses examining the effects of multiple psychiatric disorders on the cortex, more broadly defined (**Figure 6**). Across diagnoses, this relationship appeared to be primarily driven by a small group of downregulated IEGs, such that the strong correlations disappeared when using non-parametric methods. This included effects of Alcohol Abuse Disorder (AAD, *n*=32, **Figure 6A**), MDD (*n*=176, **Figure 6B**), Bipolar Disorder (BPD, *n*=197, **Figure 6C**), and Schizophrenia (SCHIZ, *n*=314, **Figure 6E**) in the cortex as measured by microarray ((Gandal et al., 2018), *MDD: R*=0.32, *p*=0.0011, *Rho*=0.04, *p*=0.702, 20 of 102 genes *p*<0.05; *BPD: R*=0.22, *p*=0.0655*\*trend*, *Rho*=-0.074, *p*=0.531, 26 of 74 genes *p*<0.05; *SCHIZ*: *R*=0.40, *p*=0.000363; *Rho*=0.11, *p*=0.314; 35 of 74 genes *p*<0.05; *AAD: R*=0.21, *p*=0.0368, *Rho*=0.15, *p*=0.142, 18 of 98 genes *p*<0.05), and effects of BPD (*n*=171, **Figure 6D**) and SCHIZ (*n*=384, **Figure 6F**) on the frontal cortex as measured by RNA-Seq ((Gandal et al., 2018), *BPD: R*=0.58, *p*=7.60e-09; *Rho*=0.11, *p*=0.328; 15 of 85 genes *p*<0.05; *SCHIZ: R*=0.37, *p*=0.000505; *Rho*=0.033, *p*=0.766; 23 of 85 genes *p*<0.05). In contrast, the DEGs driving the correlations between our meta-analysis results and other animal chronic stress studies, and similarities with PTSD and Suicide effects in the PFC, seemed to more closely reflect glial dysregulation (**Table S3**).

**Figure 6.**
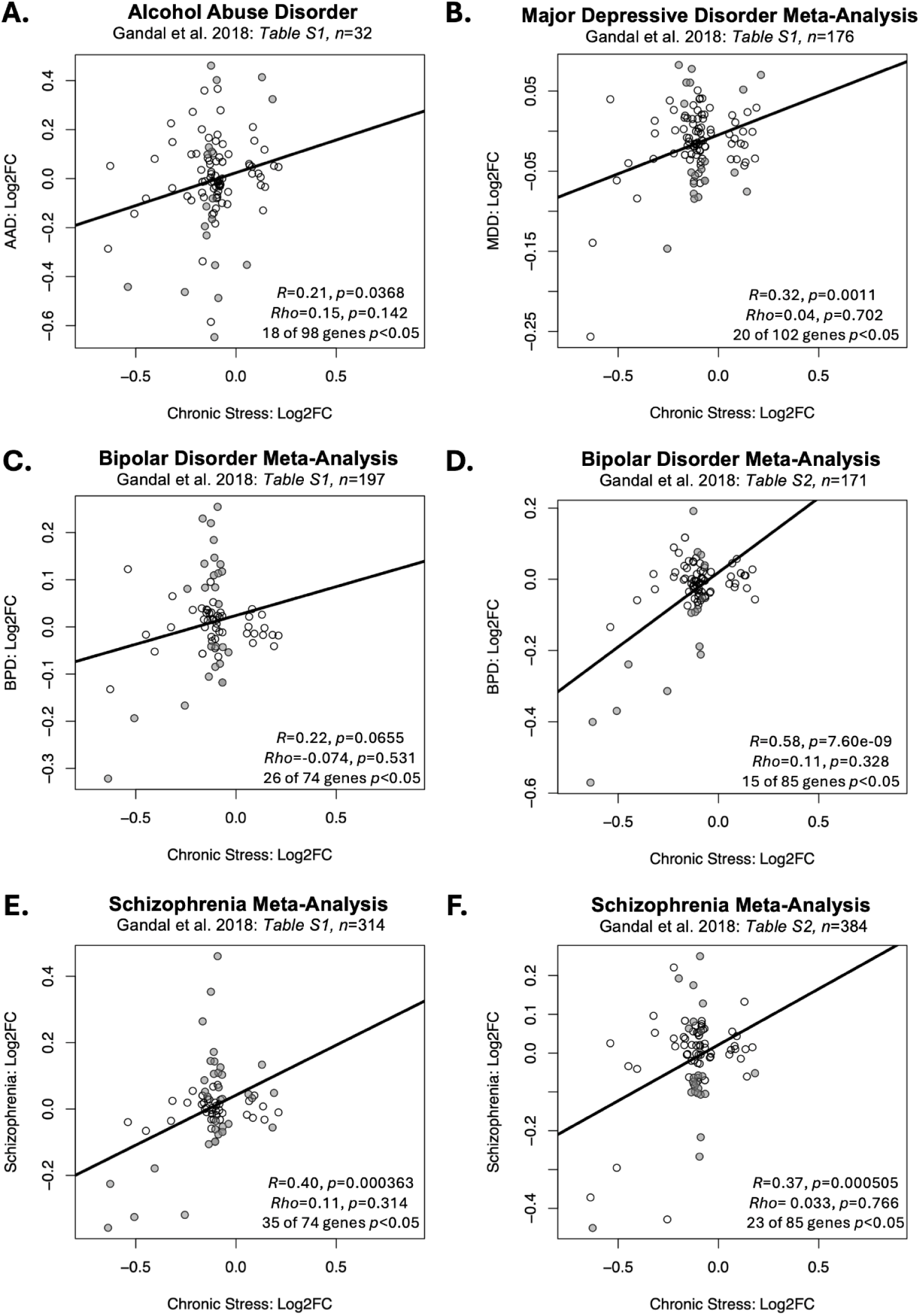
Some of the DEGs from our current meta-analysis show similar effects in previous large studies characterizing the effects of psychiatric disorders on the cortical transcriptome. **A-F)** Scatterplots illustrate the correlation between the effects of chronic stress within our meta-analysis (Log2FC: x-axis) and the effects of multiple psychiatric disorders in previous studies (Log2FC: y-axis) for any of our 133 identified DEGs (FDR<0.05) that had results available in the referenced publication. Data points are shaded grey if the effects within the previous publication were nominally significant (p<0.05). Trend lines illustrate the relationship between the two sets of results as identified by simple linear regression. Across diagnoses, these positive findings appeared to be primarily driven by a small group of downregulated IEGs, such that the strong correlations typically disappeared when using non-parametric methods. **A-C, E)** These scatterplots illustrate the relationship between the effects of chronic stress within our meta-analysis and the effects of each psychiatric disorder on the cortical transcriptome as measured by microarray (Gandal et al., 2018). Note that the Bipolar Disorder and Schizophrenia meta-analyses share overlapping control subjects and thus cannot be considered fully independent. **D, F)** These scatterplots illustrate the relationship between the effects of chronic stress within our meta-analysis and the effects of each psychiatric disorder on the frontal cortical transcriptome as measured by RNA-Seq (Gandal et al., 2018). Note that the Bipolar Disorder and Schizophrenia meta-analyses share overlapping control subjects and thus cannot be considered fully independent. **A)** Alcohol Abuse Disorder (n=32: R=0.21, Rho=0.15), **B)** MDD (n=176: R=0.32, Rho=0.04), **C)** Bipolar Disorder (n=197: R=0.22, Rho=-0.074), **D)** Bipolar Disorder (n=171: R=0.58, Rho=0.11), **E)** Schizophrenia (n=314: R=0.40, Rho=0.11), **F)** Schizophrenia (n=384: R=0.37, Rho= 0.033).

## Discussion

Our results documenting the effects of chronic stress on prefrontal gene expression add to growing evidence that the PFC remains plastic and vulnerable to stress into adulthood (Algaidi, 2025). To identify effects of chronic stress that were consistent across laboratories and models, we performed a meta-analysis of six independent mouse studies using chronic social defeat (CSDS) and chronic unpredictable stress (CUMS) paradigms (total *n*=117). Across studies 97 transcripts were consistently downregulated and 36 upregulated. The downregulated genes were highly enriched for markers of glial cells (especially astrocytes and oligodendrocytes), neurovasculature, myelination, and inflammatory or immune processes, suggesting that chronic stress broadly disrupts glial and vascular support functions in the PFC in a manner that seems to parallel previous observations from the PFC of patients with stress-related disorders such as PTSD and suicide. We also observed consistent down-regulation of IEGs across multiple datasets, paralleling what has been observed in broader cortical dissections from patients with multiple human neuropsychiatric disorders, including those that are traditionally considered stress-related (MDD) as well as those that are accompanied and exacerbated by chronic stress (AAD, BPD, SCHIZ).

### Astrocyte and Neurovascular Function

Our meta-analysis revealed that chronic stress causes consistent down-regulation of gene sets associated with astrocytic function and neurovascular integrity in the PFC, with *Aqp4*, *S100b,* and *Hepacam* amongst the leading contributors. Evidence from both clinical and preclinical studies supports the down-regulation of these genes under chronic stress.

*Aqp4* encodes aquaporin-4, a water channel protein enriched in astrocytic endfeet that is related to blood-brain barrier water homeostasis and glymphatic clearance (Mader & Brimberg, 2019, Papadopoulos & Verkman, 2013). Similar to our results, *Aqp4* was downregulated in the anterior cortex of mice following CUMS (Wei et al., 2019) and in the hippocampus following chronic corticosterone (Juszczak et al., 2025). This finding may be clinically relevant, as *AQP4* expression is reduced in the post-mortem orbitofrontal cortex (Rajkowska et al., 2013) and hippocampus (Medina et al., 2016) of patients with MDD, and in the PFC of subjects who died of suicide (Piras et al., 2022).

Gene expression related to astrocyte development was downregulated. *S100b* is an astrocyte-derived calcium binding protein that promotes glial and neuronal proliferation, survival, and differentiation (Hernández-Ortega et al., 2024). Similar to our results, CUMS reduced *S100b* mRNA and associated protein in the PFC (Bollinger et al., 2025; Luo et al., 2010), resembling the down-regulation produced by chronic corticosterone (Juszczak et al., 2025). *Hepacam* encodes an astroglial adhesion molecule regulating astrocyte territory and morphological complexity (Baldwin et al., 2021). *Hepacam* expression in the PFC has not been previously linked to chronic stress. However, *Hepacam* is sensitive to glucocorticoids (Juszczak & Stankiewicz, 2018) and *HEPACAM* shows a gene-based association with suicidality, moderated by environmental factors (Wendt et al., 2021). The down-regulation of *S100b* and *Hepacam* we observed may indicate stress-induced astrocytic structural remodeling.

Disrupted astrocytic and neurovascular gene expression following chronic stress may underlie impaired astrocyte network function. Prior work shows that chronic stress induces astrocytic atrophy without reducing cell number (Tynan et al., 2013), weakening astrocyte syncytial networks (Aten et al., 2023), dysregulating astrocyte-neuron metabolic coupling (Descalzi, 2020); and altering blood-brain barrier integrity (Dudek et al., 2020; Menard et al., 2017). These mechanisms are increasingly thought to be related to mood disorders (e.g, (Nestler & Russo, 2024)).

### Oligodendrocytes

Chronic stress produced down-regulation in gene sets representing mature oligodendrocytes, oligodendrocyte precursor cells, and myelination processes in the PFC. This included down-regulation of two enzymes, *Fa2h* and *Elovl5,* that are responsible for the synthesis of critical myelin components (Hoxha et al., 2021; Potter et al., 2011). Consistent with our results, *Fa2h* was downregulated in the hippocampus following chronic corticosterone (Juszczak et al., 2025), and in the PFC following CSDS (Cathomas et al., 2019) in stress susceptible animals (Reshetnikov et al., 2022), and also potentially after early life stress (Duan et al., 2025). *Elovl5* was also down-regulated following chronic corticosterone (Juszczak et al., 2025). These findings could be clinically important, as *Fa2h* was decreased in patients with PTSD (Stankiewicz et al., 2022a), and *ELOVL5* is reduced in the postmortem frontal cortex in patients with MDD (McNamara & Liu, 2011), alcohol abuse disorder (Gandal et al., 2018), and suicide (Piras et al., 2022). These molecular changes align with a broader pattern of oligodendrocyte and myelination abnormalities reported in the PFC of stressed animals (Laine et al., 2018; Lehmann et al., 2017) and patients with stress-related disorders, including MDD and BD (Czéh & Nagy, 2018).

### Glial Communication

Chronic stress suppressed genes related to glial communication, including *P2rx7*. *P2rx7* encodes a purinergic receptor, P2X7R, that is activated by stress-elevated extracellular ATP, triggering downstream neuroinflammation linked to depression (Adinolfi et al., 2018; von Muecke-Heim et al., 2021). In chronic stress models, P2X7R antagonists consistently blunted depressive-like behaviors and reversed immune changes (Ribeiro et al., 2019). *P2rx7* down-regulation in our meta-analysis and in patients with MDD (Aston et al., 2005) might reflect a homeostatic or compensatory response to chronic overstimulation of this pathway, limiting excessive inflammation. Overall, the down-regulation of multiple glial-driven gene sets suggests that chronic stress may alter glial-neuron communication and neuroimmune signaling within PFC circuits.

### Immediate Early Genes

Chronic stress suppressed immediate early genes (IEGs) linked to neuronal activity and plasticity, such as *Fos*, *Junb*, *Arc*, *Egr1,* and *Dusp1* (**Figure 4**) (Abate et al., 1991; Chen et al., 2019; Korb & Finkbeiner, 2011). A comprehensive review reported that these IEGs are reliably induced by acute stressors but down-regulated after chronic stress across multiple regions, including the hippocampus and PFC (Flati et al., 2020) in a manner important for social behavioral stress responses in susceptible animals (Inaba et al., 2023). However, this down-regulation of IEGs following chronic stress may be context dependent, as some studies have found the opposite pattern (*e.g.,* (Deonaraine et al., 2020; Zhao et al., 2017), **Fig S2 C&D**). As a similar down-regulation of IEG expression was observed in the cortex across a variety of psychiatric disorders, including alcohol abuse disorder, MDD, bipolar disorder, and schizophrenia (Gandal et al., 2018), this complex relationship between IEG expression and chronic stress is worth investigating further.

### Upregulation of Xdh

Finally, we observed an upregulation of *Xdh* that was consistent with nominal findings from other meta-analyses of chronic stress (Gururajan, 2022; Reshetnikov et al., 2022) and stress susceptibility (Reshetnikov et al., 2022), and large studies of chronic corticosterone (Juszczak et al., 2025) and prolonged stress (Stankiewicz et al., 2022a). Upregulation of Xdh was also associated with schizophrenia (Gandal et al., 2018) (**Table S3**). *Xdh* encodes the protein xanthine oxidoreductase (XOR), the terminal enzyme of purine catabolism (Furuhashi, 2020). Chronic stress increases cortical XOR activity, resulting in cerebrovascular dysfunction and cognitive deficits (Burrage et al., 2023). Upregulation of *Xdh* following chronic stress exposure may similarly promote a shift towards increased oxidative stress at the neurovascular interface.

### Strengths and Limitations

Compared to the original studies, our meta-analysis provides greater statistical power, allowing more robust identification of DEGs, and better generalizability of findings. However, our analysis has several limitations.

First, the limited number of comparisons between stress susceptible and resilient groups did not allow us to properly examine how stress susceptibility affects PFC gene expression. However, a recent meta-analysis of CSDS effects on the PFC did pursue this question (Reshetnikov et al., 2022). When compared with our meta-analysis, their results suggest that susceptible animals may have generalized stress effects that are amplified compared to resilient animals, such as greater down-regulation of myelin-related genes (Reshetnikov et al., 2022). These findings are an interesting complement to previous work, which revealed active compensatory mechanisms bolstering stress resilience (Nestler & Russo, 2024).

Second, behavioral responses to chronic stress have been shown to be sex-specific (e.g., (Hodes et al., 2015)). Since our datasets were derived predominantly from male mice, our findings may be limited to males. To explore this possibility, we conducted a small follow-up analysis comparing our meta-analysis results to published differential expression results broken down by sex (Deonaraine et al., 2020; Labonté et al., 2017; Shao et al., 2025). These comparisons suggested that the chronic stress DEGs identified by our meta-analysis show similar effects in males and females (**Figure S2**). However, these conclusions are based on limited published results from small studies and should be followed up with a meta-analysis designed to characterize sex differences.

Finally, our findings reflect samples encompassing a variety of prefrontal cortical regions. These regions are likely to serve different roles in stress responses and exhibit prominent variation across species (Preuss & Wise, 2022). They also include a variety of cell types with divergent stress responses (Kwon et al., 2022; Shao et al., 2025) which may not be fully reflected in our results derived from bulk dissections. Future research will be necessary to explore the regional and cell type specificity of our identified effects.

### Conclusion

Chronic stress is a significant public health concern in the United States, contributing to the onset and exacerbation of psychiatric conditions (Davis et al., 2017). Human studies often have modest effect sizes and complex designs, making it difficult to identify targets. By releasing our full meta-analysis results **(Table S1**: 21,379 genes, with 133 DEGs) and gene set enrichment results (**Table S2**: 10,847 gene sets, 53 with FDR<0.05), we provide a useful reference database that can be compared to the transcriptional profiling results from psychiatric and neurological conditions to distinguish stress-associated factors. Moreover, our convergent analyses with prior human transcriptomic reports highlighted patterns that are consistent across species, providing insight for mechanistic and translational follow-up.

## Supporting information

Supplemental Methods & Results

## CRediT Statement

JX: Conceptualization, Methodology, Software, Formal analysis, Investigation, Data Curation, Writing - Original Draft, Writing - Review & Editing, Visualization

MHH: Conceptualization, Methodology, Software, Formal analysis, Writing - Original Draft, Writing - Review & Editing, Validation, Visualization, Supervision, Project administration

CAR: Methodology, Writing - Review & Editing

NRC: Writing - Review & Editing

EIF: Methodology, Writing - Review & Editing

EH: Methodology, Writing - Review & Editing

DMN: Methodology, Writing - Review & Editing

AS: Methodology, Writing - Review & Editing

TQD: Writing - Review & Editing

SJW: Writing - Review & Editing, Funding acquisition

HA: Writing - Review & Editing, Supervision, Funding acquisition

## Acknowledgements

This work was completed as part of the *Brain Data Alchemy Project* and supported by the Hope for Depression Research Foundation (HDRF: HA), Grinnell College Center for Careers, Life, and Service (JX, CAR, EH, DMN, TQD), the International Brain Research Organization (IBRO) and Faculty for Undergraduate Neuroscience (FUN) (DMN, TQD), National Institute on Drug Abuse (NIDA U01 DA043098: HA & SJW), and the Pritzker Neuropsychiatric Disorders Research Foundation (HA & SJW). Funders and sponsors had no active role in the review.

## Data Availability Statement

All datasets used in this publication are publicly available in the National Center for Biotechnology Information (NCBI) repository Gene Expression Omnibus (GEO, https://www.ncbi.nlm.nih.gov/geo/) under the accession numbers GSE84572, GSE109315, GSE81672, GSE81587, GSE102556, and GSE151807.

