## Supplemental Methods & Results for "A Meta-Analysis of the Effects of Chronic Stress on the Prefrontal Transcriptome in Animal Models and Convergence with Existing Human Data"

*Shared first authorship

Direct correspondence to:

Megan Hagenauer, Ph.D.
Michigan Neuroscience Institute
BSRB
109 Zina Pitcher Pl
University of Michigan
Ann Arbor, MI 48109 USA
**

### Supplemental Methods

#### *Comparison of Meta-Analysis Results to Large Studies Characterizing Acute Stress Effects on the Brain:*

We compared our meta-analysis results to several studies examining acute stress effects on the brain using either meta-analyses or large sample sizes. First, the comprehensive database included in the supplement of (Juszczak et al., 2025: *Suppl Data 2)* included the full summarized results from a large study of acute corticosterone exposure (12 hr) on the hippocampal transcriptome (Jaszczyk et al., 2023: n=48), and the results of large *"vote-counting"*-style reviews of the literature characterizing the effects of glucocorticoids (Juszczak & Stankiewicz, 2018), and acute and chronic stress on the brain transcriptome (Stankiewicz et al., 2022: mice (58 studies), rats (20 studies), squirrel monkeys (1 study)). The summarized results in this database provided categorical information about the direction of effect (up, down, mixed effects) for genes that had significant effects in the original studies.

We also compared our result to two large studies examining the effects of acute sleep deprivation on the cortical transcriptome. The first study was a large meta-analysis (Rhoads et al. 2025, *n*=293) with the full results available for all genes (*Table S1).* The second study represented pooled samples from a large sample size (Diessler et al. 2018, GSE114845: n=86 RNA-Seq samples from n=222 mice), with the full results for all genes available following re-analysis in (Rhoads et al. 2025: *Table S2*).

***Statistical Methods for Result Comparisons:*** To conduct these comparisons, we extracted all available results for our 133 differentially expressed genes ("DEGs" FDR<0.05). If there were Log2FC values available, we examined the correlation with the Log2FC estimated from our chronic stress meta-analysis using both parametric (regression, Pearson's correlation) and non-parametric (Spearman's correlation) methods (functions: *summary.lm, cor.test*). If only direction of effect information was available, we only compared our results for genes with a clear "Up" or "Down" direction of effect. For acute corticosterone exposure (12 hr) (Jaszczyk et al., 2023), there were three collection timepoints - we allowed results that showed "Up" or "Down" regulation at any of the three timepoints, but excluded genes that showed effects in different directions at different timepoints. The relationship between previous documented direction of effect and the Log2FC estimated from our chronic stress meta-analysis was then analyzed using a Welch's two-tailed t-test (function: *t.test*).

#### *Comparison of Meta-Analysis Results to Published Effects of Chronic Stress in Each Sex:*

Behavioral responses to chronic stress have been previously shown to be sex-specific (e.g., (Hodes et al., 2015)). One limitation of our meta-analysis is that our sample was overwhelmingly male. To explore whether our meta-analysis findings might be male-specific, we conducted a small follow-up analysis comparing our meta-analysis results to published differential expression results broken down by sex (Deonaraine et al., 2020; Labonté et al., 2017; Shao et al., 2025). These comparisons included a CUMS study that was part of our meta-analysis that included both males and females ((Labonté et al., 2017): *n*=19 males and *n*=19 females). For our meta-analysis, we used the chronic stress vs. control effect estimated from the full sample (both sexes: *n*=38 of our full sample size of *n*=117). However, the original publication included differential expression results for *chronic stress vs. control* comparisons for all genes that survived a nominal $p<0.05$ threshold in the male PFC (*Table S17:* 1412 genes) and female PFC (*Table S18:* 2132 genes), analyzed separately, allowing us to explore the sex specificity of our observed effects. A recent publication also characterized the effects of CUMS in each sex ((Shao et al., 2025), *n*=11 males and *n*=10 females) in their genetic intervention control (*SSTCre*) condition. They provided the differential expression results for *chronic stress vs. control* comparisons for all genes that survived a nominal $p<0.01$ threshold in the male PFC (*Table S1*: 437 genes) and female PFC (*Table S7*: 402 genes), analyzed separately. We also compared our meta-analysis findings to a CSDS experiment that was excluded during our inclusion/exclusion procedure (Deonaraine et al., 2020). CSDS is traditionally a male-only chronic stress paradigm, because mice typically only show territorial aggression towards intruder males. This study modified the CSDS paradigm to function as a chronic stress paradigm for females by using genetically-modified aggressor mice that would act aggressively towards females (Deonaraine et al., 2020). They then compared their findings to males from a previously published CSDS experiment from their group (Wang et al., 2018). They provided differential expression results for chronic stress vs. control comparisons for the vehicle-treated mice (*n*=9 male samples, *n*=8 female samples (pooled from 3-4 mice) for all genes that survived a p<0.05 threshold in the females (*Table S1*: 1271 genes) and males (*Table S3:* 2698 genes), analyzed separately.

To compare the published results to our meta-analysis results, we extracted all available results for our 133 differentially expressed genes ("DEGs" FDR<0.05) from each of the publications. We then examined the correlation with the Log2FC estimated from our chronic stress meta-analysis using both parametric (regression, Pearson's correlation) and non-parametric (Spearman's correlation) methods (functions: *summary.lm, cor.test*).

### Supplemental Results

#### *Comparison of Meta-Analysis Results to Large Studies Characterizing Acute Stress Effects on the Brain:*

When comparing our chronic stress meta-analysis results to the effects of acute daily corticosterone on the hippocampal transcriptome (Jaszczyk et al., 2023: *n*=48 subjects), there did appear to be some similarities, with DEGs that were previously upregulated in response to acute corticosterone (n=29 genes) tending to have greater Log2FCs in our chronic stress meta-analysis than those that were previously down-regulated in response to acute corticosterone (n=26 genes), but this relationship was not significant (T(51.36)=-1.45, p=0.153).

Similarly, when comparing our findings to a large meta-analysis of the effects of acute sleep deprivation on the cortical transcriptome (Rhoads et al. 2025, *n*=293 subjects), results were available for 118 of our DEGs, of which 17 showed nominal significance (p<0.05). The Log2FCs for these genes showed a nice positive correlation with the Log2FCs in our meta-analysis (118 genes: R=0.70, p<2.2e-16, Rho=0.37, p=4.27e-05, **Fig 5F**). However, the opposite pattern was observed when comparing our meta-analysis results to the effects of acute sleep deprivation measured in a separate large study (Diessler et al. 2018, GSE114845: n=86 RNA-Seq samples from n=222 mice). For this study, results were available for 116 of our DEGs, for which 81 showed nominal significance (p<0.05). The Log2FCs for these genes showed a negative correlation with the Log2FCs in our meta-analysis, but this negative correlation appeared to be driven by a handful of IEGs, so the negative correlation disappeared when running a non-parametric analysis (R=-0.45, p=2.92e-07, Rho=-0.017, p=0.851).

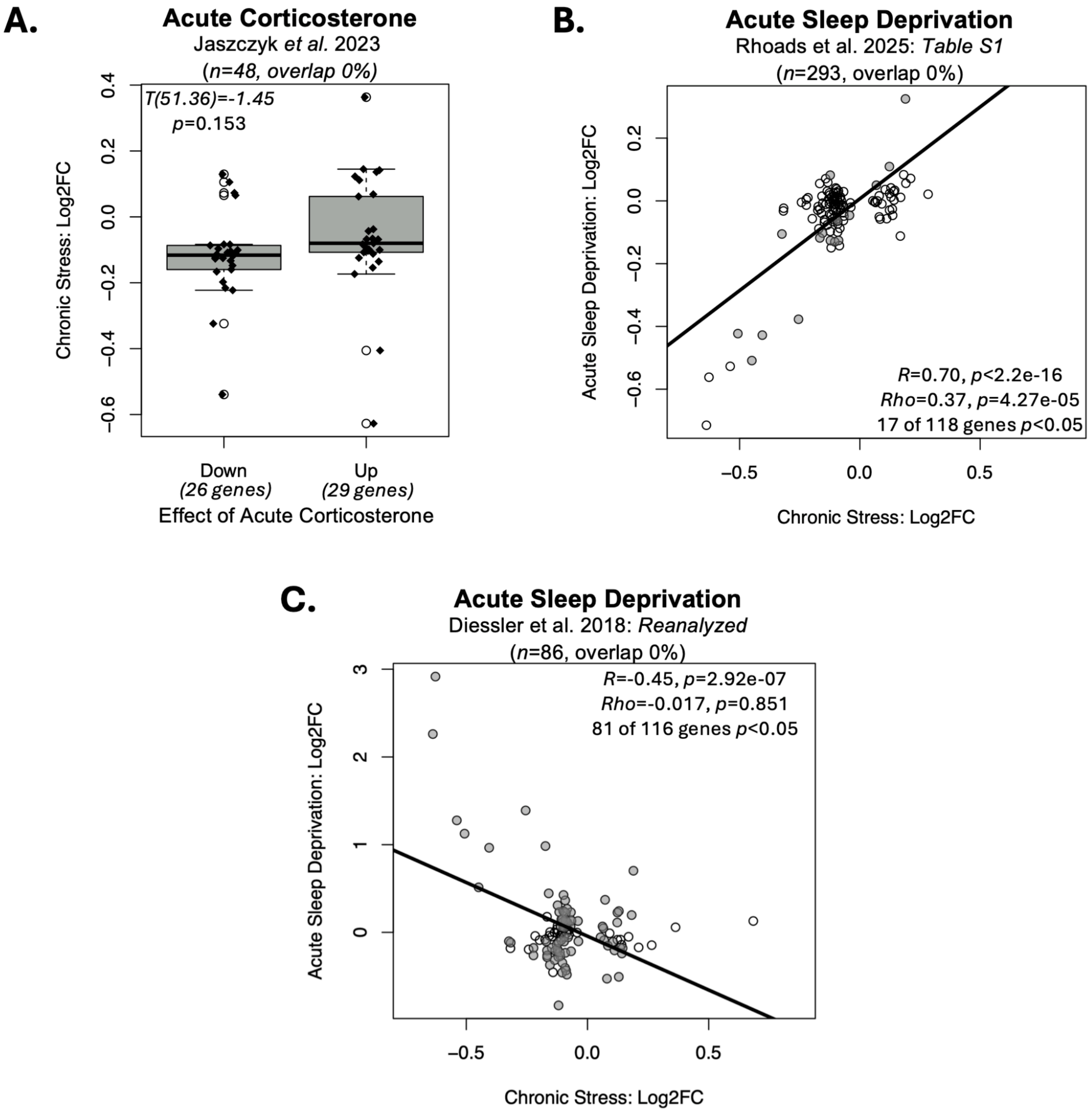


**Figure S1.** ***The DEGs from our current chronic stress meta-analysis often show effects in previous large studies characterizing the effects of acute stress and glucocorticoids on the brain transcriptome. A.*** *Within a previous large study examining the effects of acute daily corticosterone on the hippocampal transcriptome (Jaszczyk et al., 2023: n=48 subjects), 55 of our DEGs showed consistent significant differential expression. For these genes, a boxplot*

*illustrates the relationship between the direction of effect of acute daily corticosterone (upregulated: 29 genes, down-regulated: 26 genes) and the Log2FCs in our chronic stress meta-analysis. This relationship was not significant (T(51.36)=-1.45, p=0.153).* ***B-C:*** *Scatterplots illustrate the correlation between the effects of chronic stress within our meta-analysis (Log2FC: x-axis) and the effects of stress within previous studies (Log2FC: y-axis) for any of our 133 identified DEGs (FDR<0.05) that had results available in the referenced publication. Data points are shaded grey if the effects within the previous publication were nominally significant (p<0.05). Trend lines illustrate the relationship between the two sets of results as identified by simple linear regression.* ***B.*** *Differential expression results were available for 118 of our DEGs within a large meta-analysis of the effects of acute sleep deprivation on the cortical transcriptome (Rhoads et al. 2025; n=293 subjects), For these genes, there was a positive correlation with the Log2FCs in our meta-analysis (118 genes: R=0.70, Rho=0.37).* ***C.*** *Differential expression results were available for 116 of our DEGs within a large study examining the effects of sleep deprivation on the cortical transcriptome (Diessler et al. 2018, GSE114845: n=86 RNA-Seq samples from n=222 mice). For these genes, there was a negative correlation with the Log2FCs in our meta-analysis (R=-0.45, Rho=-0.017), but this negative correlation appeared to be driven by a handful of IEGs, so that the negative correlation disappeared when running a non-parametric analysis.*

***Comparison of Meta-Analysis Results to Published Effects of Chronic Stress in Each Sex:***

Response to chronic stress has been previously shown to be sex-specific. For example, six days of variable stress could induce depression-associated behaviors in female mice but not in males (Hodes et al., 2015). One limitation of our meta-analysis is that our sample was overwhelmingly male. To explore whether our meta-analysis findings might be male-specific, we conducted a small follow-up analysis comparing our meta-analysis results to published differential expression results broken down by sex (Deonaraine et al., 2020; Labonté et al., 2017; Shao et al., 2025). These comparisons included a CUMS study that was part of our meta-analysis that included both males and females ((Labonté et al., 2017): $n$=19 males and $n$=19 females). For our meta-analysis, we used the chronic stress vs. control effect estimated from the full sample (both sexes: $n$=38 of our full sample size of $n$=117). However, the original publication included differential expression results for *chronic stress vs. control* comparisons for all genes that survived a nominal $p<0.05$ threshold in the male PFC (1412 genes) and female PFC (2132 genes) analyzed separately. These reported results included male-derived differential expression results for 16 of the chronic stress DEGs identified by our meta-analysis and female-derived differential expression results for 14 of our DEGs. For these genes, in both males and females there was a strong positive correlation with the Log2FCs from our chronic stress meta-analysis (**Figure S2A:** ***Males:*** 16 DEGs: $R$=0.96, $p$=1.78e-09, $Rho$=0.85, $p$=2.02e-06; ***Females:*** 14 DEGs: $R$=0.82, $p$=0.000284, $Rho$=0.76, $p$=0.00231).

We observed a similar pattern when comparing our meta-analysis results to findings from a recent publication characterizing the effects of CUMS in each sex ((Shao et al., 2025), $n$=11 males and $n$=10 females) in their genetic intervention control (*SSTCre*) condition. They provided the differential expression results for *chronic stress vs. control* comparisons for all genes that survived a nominal $p<0.01$ threshold in the male PFC (437 genes) and female PFC (402 genes), analyzed separately. These reported results included male-derived differential expression results for 10 of the chronic stress DEGs identified by our meta-analysis and female-derived differential expression results for 4 of our DEGs. For these genes, in both males and females there was a strong positive correlation with the Log2FCs from our chronic stress meta-analysis (**Figure S2B:** ***Males:*** 10 DEGs: $R$=0.86, $p$=0.00151, $Rho$=0.92, $p$=0.000467; ***Females:*** 4 DEGs: $R$=0.95, $p$=0.0539, $Rho$=1, $p$=0.0833), although the findings from the females should be taken tentatively due to the small number of genes included in the comparison and weaker p-value (non-significant trend).

We also compared our meta-analysis findings to a CSDS experiment that was excluded during our inclusion/exclusion procedure (Deonaraine et al., 2020). This study modified the CSDS paradigm to function as a chronic stress paradigm for females by using genetically-modified aggressor mice that would act aggressively towards females (Deonaraine et al., 2020). They then compared their findings to males from a previously published CSDS experiment from their group (Wang et al., 2018). They provided differential expression results for chronic stress vs. control comparisons for the vehicle-treated mice ($n$=9 males, $n$=8 females) for all genes that survived a $p<0.05$ threshold in the females (1271 genes) and males (2698 genes), analyzed separately. These reported results included male-derived differential expression results for 25 of the chronic stress DEGs identified by our meta-analysis and female-derived differential expression results for 12 of our DEGs. For these genes, both males and females showed a negative correlation with the Log2FCs from our chronic stress meta-analysis (**Figure S2C**: ***Males:*** 25 DEGs: $R$=-0.47, $p$=0.01765, $Rho$=-0.25, $p$=0.236; ***Females:*** *12 DEGs: R*=-0.66, *p*=0.0184, *Rho*=-0.64, *p*=0.0280), in a manner that seemed to be at least partially driven by a large upregulation of immediate early genes in both sexes (**Figure S2D**). These findings resemble the upregulation of immediate early genes that is sometimes observed following acute stress (e.g., **Fig S2C**, (Flati et al., 2020), but the brain samples were collected three days following CSDS and two days following behavioral testing (Deonaraine et al., 2020). Therefore, these findings seem to provide an interesting counterexample to our meta-analysis findings, but do not provide evidence that the chronic stress effects that we detected are sex-specific.

Based on these comparisons, we tentatively conclude that the general chronic stress DEGs identified by our meta-analysis do not appear to show sex-specific differential expression when the sexes are analyzed independently. However, these conclusions should be considered weak, at best, since all of the published results included in our comparisons were thresholded by statistical significance within the original studies. Therefore, our comparisons are only able to identify effects that showed a significant *reversal* in females. We are unable to make a strong statement regarding genes that may *lack* an effect in females that was detected in our meta-analysis, especially since false negative results are prevalent in transcriptional profiling studies using small samples and many of the male-only studies included in our meta-analysis also did not show significant effects for these genes either when analyzed independently (*e.g.,* examples in the forest plots provided in **Fig 3** & **Fig 4**). Future well-powered studies will be needed to better address the question, such as a meta-analysis of sex differences in the effects of chronic stress.

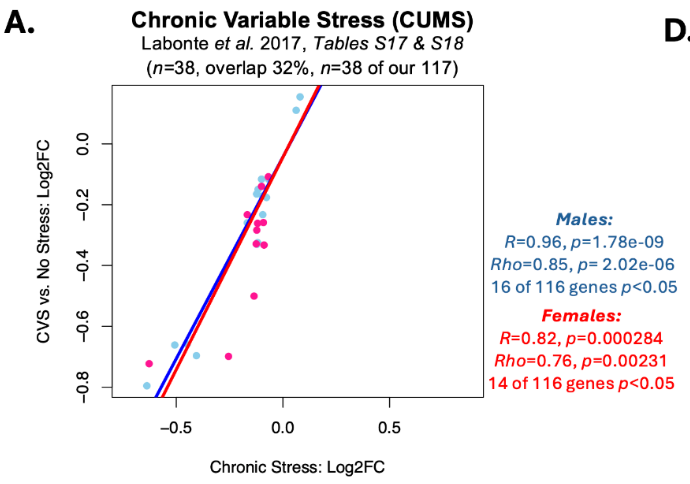


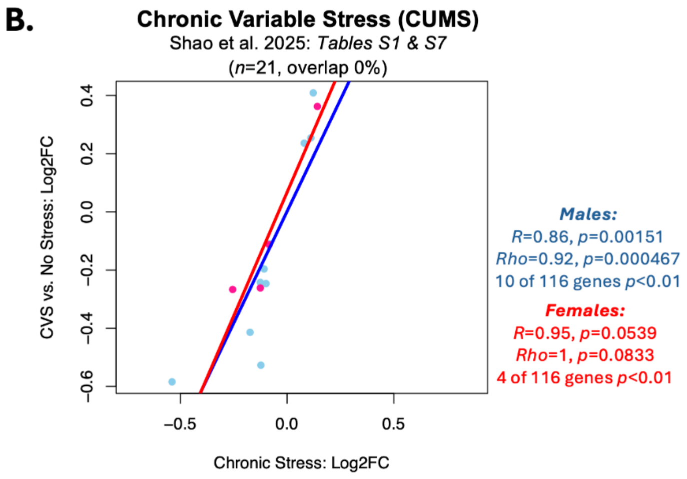


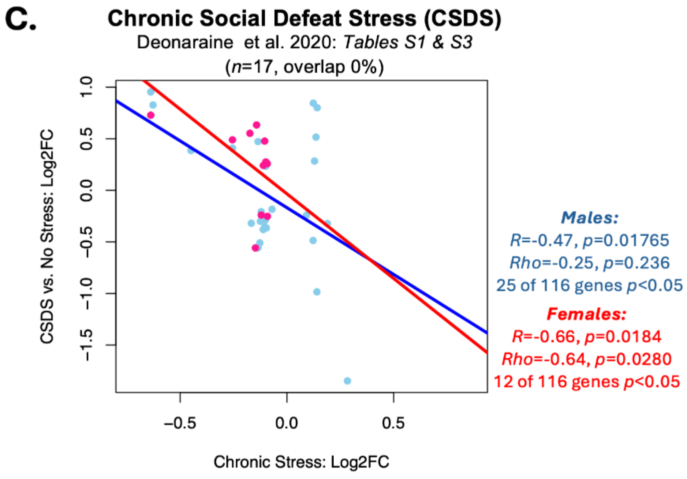


**D.**

| GeneSymbol | ChronicStress_Xiong_Log2FC_estimate | Labonte_Male_CVSvsNS_logFC | Labonte_Female_CVSvsNS_logFC | Shao_Male_CVSvsNS_SSTCre_log2FoldChange | Shao_Female_CVSvsNS_SSTCre_log2FoldChang | Deoraine_2020_male_CSDSvsNoStress_logFC | Deoraine_2020_female_CSDSvsNoStress_logFC |
|---|---|---|---|---|---|---|---|
| *Fos* | -0.64 | -0.80 | -0.91 | | | 0.95 | 0.73 |
| *Arc* | -0.63 | | -0.72 | | | 0.83 | 1.25 |
| *Ccn1* | -0.54 | | | -0.58 | | | |
| *Dusp1* | -0.51 | -0.66 | | | | | |
| *Btg2* | -0.45 | -0.66 | | | | 0.38 | |
| *Ier2* | -0.41 | -0.70 | | | | | |
| *Egr1* | -0.26 | | -0.70 | | -0.27 | 0.41 | 0.49 |
| *Csrnp1* | -0.17 | | | -0.41 | | | |
| *Fam163a* | -0.17 | -0.26 | -0.23 | | | -0.32 | |
| *Atad2* | -0.15 | | | | | | -0.56 |
| *Tnni1* | -0.14 | | | | | | 0.63 |
| *Syndig1* | -0.14 | | | | | -0.56 | |
| *Cd274* | -0.14 | | -0.50 | | | 0.47 | |
| *S100b* | -0.13 | | | | | -0.51 | |
| *Aqp4* | -0.13 | | -0.33 | -0.24 | -0.26 | | |
| *Lgi3* | -0.12 | | | | | -0.30 | |
| *Ppp1r16b* | -0.12 | -0.16 | | | | | |
| *Kdr* | -0.12 | -0.33 | -0.28 | -0.53 | | | |
| *Rbms1* | -0.12 | | | | | -0.21 | -0.24 |
| *Tek* | -0.12 | -0.33 | -0.26 | | | | |
| *Siae* | -0.12 | -0.15 | | | | | |
| *Mxra7* | -0.11 | | 0.24 | | | | |
| *Adgra3* | -0.11 | | | | | -0.38 | |
| *Gna12* | -0.11 | | | | | | 0.24 |
| *Elovl5* | -0.11 | | | -0.20 | | | |
| *Slc39a8* | -0.1 | | | | | | 0.48 |
| *Rnd2* | -0.10 | | | | | -0.35 | |
| *Anxa5* | -0.10 | | | | | -0.29 | |
| *Atf4* | -0.10 | -0.12 | -0.14 | | | | |
| *Smad7* | -0.10 | | | -0.25 | | 0.23 | 0.28 |
| *Car10* | -0.10 | | | | | -0.36 | |
| *Map3k11* | -0.10 | -0.17 | | | | | |
| *Rgs8* | -0.09 | -0.23 | | | | | |
| *Bag3* | -0.09 | | | | | | 0.26 |
| *Zfp362* | -0.09 | | -0.26 | | | | -0.25 |
| *Alpl* | -0.09 | | -0.33 | | | | |
| *Trp53inp2* | -0.08 | | | -0.11 | -0.11 | | |
| *Kif1c* | -0.08 | -0.18 | | | | | |
| *Bckdhb* | -0.07 | | | | | -0.18 | |
| *Sec61a1* | -0.07 | | -0.11 | | | | |
| *Icmt* | -0.07 | -0.12 | | | | | |
| *Scaf8* | 0.06 | 0.11 | | | | | |
| *Phka2* | 0.08 | 0.15 | | 0.24 | | | |
| *Sidt1* | 0.09 | | | | | -0.24 | |
| *Slc24a4* | 0.11 | | | 0.25 | | | |
| *Nrros* | 0.12 | | | 0.41 | | -0.49 | |
| *Myo1h* | 0.12 | | | | | 0.84 | |
| *Psrc1* | 0.13 | | | | | 0.28 | |
| *Egfem1* | 0.14 | | | | | 0.52 | |
| *Qprt* | 0.14 | | | | | -0.98 | |
| *Col6a2* | 0.14 | | | | 0.36 | 0.80 | |
| *Cox4i2* | 0.14 | | 0.43 | | | | |
| *Xdh* | 0.19 | | | | | -0.32 | |
| *Nphs1os* | 0.28 | | | | | -1.85 | |

***Figure S2. The DEGs identified by our chronic stress meta-analysis do not appear to show effects that notably differ by sex in previously published sex-specific analyses. A.*** *For one of* *the studies included in our meta-analysis (Labonté et al., 2017) the effects of chronic stress were provided in the original publication for males (n=19) and females (n=19), analyzed separately (only nominally significant p<0.05 results available). Differential expression results from the males were available for 16 of our DEGs, and from the females there were results available for 14 of our DEGs. For these genes, both sexes showed effects (Log2FCs) that had a strong positive correlation with the Log2FCs from our meta-analysis (males: R=0.96, Rho=0.85, females: R=0.82, Rho=0.76).* ***B.*** *A recent publication characterized the effects of CUMS in each sex ((Shao et al., 2025), n=11 males and n=10 females) in their genetic intervention control (SSTCre) condition (only nominally significant p<0.01 results available). Differential expression results from the males were available for 10 of our DEGs, and from the females there were results available for 4 of our DEGs. For these genes, both sexes showed effects (Log2FCs) that had a strong positive correlation with the Log2FCs from our meta-analysis (males: R=0.86, Rho=0.92; females: R=0.95, Rho=1), although the findings from the females should be taken tentatively due to the small number of genes included in the comparison.* ***C.*** *A study that was excluded during our inclusion/exclusion procedure (Deonaraine et al., 2020). used a modified protocol to examine the effects of CSDS in female mice and then compared their findings to males from a previously published CSDS experiment from their group (Wang et al., 2018). They provided differential expression results from the chronic stress vs. control comparisons for the vehicle-treated mice from both sexes (n=9 males, n=8 females, only nominally significant p<0.01 results available). Differential expression results from the males were available for 25 of our DEGs, and from the females there were results available for 12 of our DEGs. For these genes, both sexes showed effects (Log2FCs) that had a negative correlation with the Log2FCs from our chronic stress meta-analysis (males: R=-0.47, Rho=-0.25; females: R=-0.66, Rho=-0.64).* ***D.*** *A table providing the available Log2FCs for each sex for each of the studies visualized in panels A-C. Note the strong upregulation of immediate early genes (e.g., Fos, Arc, Egr1) in both males and females in the results from (Deonaraine et al., 2020), which contradicts the pattern of down-regulation observed in our meta-analysis.*

### Supplemental Tables and Table Legends

***Table S1. The full meta-analysis results (21,379 genes, 21,296 stable meta-analysis estimates).*** *This .xlsx file includes two worksheets: 1) The worksheet "MetaAnalysisOutputByPval" provides the full meta-analysis results, with each row representing the results for one gene, and each column providing either gene annotation or meta-analysis statistical output. The results are ordered by p-value, so that the top rows in the worksheet are the genes with the smallest p-values. 2) The worksheet "ColumnDefinitions" provides the definitions for the variables present in each column in "MetaAnalysisOutputByPval".*

***Table S2: The full fast Gene Set Enrichment Analysis (fGSEA) results (10847 gene sets).*** *This .xlsx file includes two worksheets: 1) The worksheet "fGSEA_Results" provides the full fGSEA results, with each row representing the results for one gene set, and each column providing the fGSEA statistical output. The results are ordered by p-value, so that the top rows in the* *worksheet are the gene sets with the smallest p-values. 2) The worksheet "ColumnDefinitions" provides the definitions for the variables present in each column in "fGSEA_Results".*

| GeneSymbol | ChronicStress_Xiong_Log2FC | CSDS_10Days_Reshetnikov2022_Log2FC | CSDS_30Days_Reshetnikov2022_Log2FC | SusceptibleVsResilient_Reshetnikov2022_Log2FC | ChronicStress_Gururajan2022_Log2FC | EarlyLifeStress_Duan2025_Log2FC | ChronicDailyCorticosterone_Jaszczyk2025_Log2FC | MediumLengthStress_Stankiewicz2022_Direction | ProlongedStress_Stankiewicz2022_Direction | PTSD_Stankiewicz2022_Direction | Piras2022_Suicide_AveLog2FC | Gandal_Microarray_AAD.beta_Log2FC | Gandal_Microarray_MDD.beta_Log2FC | Gandal_Microarray_SCZ.beta_Log2FC | Gandal_Microarray_BD.beta_Log2FC | Gandal_RNASeq_SCZ_Log2FC | Gandal_RNASeq_BD_Log2FC |
|---|---|---|---|---|---|---|---|---|---|---|---|---|---|---|---|---|---|
| **Consistent Effects in 2 or More Human Psychiatric Studies that Resemble Chronic Stress** | | | | | | | | | | | | | | | | | |
| ***Fos*** | -0.64 | | | | | | | Down | Mixed | | | | | -0.36 | -0.32 | | -0.57 |
| ***Arc*** | -0.63 | | | | | -0.31 | 0.48 | Down | Mixed | | | | | -0.23 | | -0.45 | -0.40 |
| ***Dusp1*** | -0.51 | -0.20 | | | | | | | Mixed | | | | | -0.33 | -0.19 | | -0.37 |
| ***Egr1*** | -0.26 | | | | | | 0.23 | | | | | -0.46 | -0.15 | -0.32 | -0.17 | | -0.31 |
| ***Syndig1*** | -0.14 | | | | | | | | | | | | -0.05 | -0.11 | -0.11 | -0.10 | -0.09 |
| ***Ppp1r16b*** | -0.12 | -0.11 | | -0.16 | | | 0.30 | | Mixed | | | | -0.07 | | -0.04 | -0.06 | |
| ***Siae*** | -0.12 | -0.10 | | | | | | Up | | | | -0.08 | -0.06 | | | -0.10 | -0.09 |
| ***Slc39a8*** | -0.10 | | | | | | -0.34 | | | | | -0.35 | | -0.10 | | -0.10 | |
| ***Rcan2*** | -0.10 | | | -0.16 | | | | | | | | | | -0.06 | -0.08 | | -0.08 |
| ***Desi1*** | -0.10 | | | | | | -0.11 | | | | | | | -0.04 | -0.04 | -0.06 | |
| ***Rgs8*** | -0.09 | | | | | | 0.12 | | Mixed | | | | | | | -0.27 | -0.19 |
| ***Alpl*** | -0.09 | | | | | | -0.68 | | | | | -0.49 | | -0.08 | | -0.22 | -0.21 |
| ***Ube2d1*** | -0.08 | | | | | | | | | | | | | | -0.08 | -0.06 | -0.05 |
| ***Pde8b*** | -0.07 | | | | | | | | | | | | -0.06 | -0.07 | -0.12 | | |
| ***Icmt*** | -0.07 | | | | | | | | | | | | | -0.05 | -0.04 | -0.11 | |
| ***Rtl8c*** | -0.04 | | | | | | | | | | | | | -0.04 | -0.05 | | |
| ***Psrc1*** | 0.13 | | | | | | -0.17 | | | | | 0.41 | | 0.13 | | | |
| **Consistent Effects in Two or More Chronic Stress Studies that Resemble At Least One Psychiatric Disorder** | | | | | | | | | | | | | | | | | |
| ***Ccn1*** | -0.54 | -0.28 | | | | | | | | | | -0.44 | | | | | |
| ***Btg2*** | -0.45 | -0.21 | | | | | 0.12 | | Mixed | | | | | | | | -0.24 |
| ***Ier2*** | -0.41 | | -0.25 | | | | | | | | | | | -0.18 | | | |
| ***Fa2h*** | -0.32 | | | -0.49 | | -0.30 | -0.19 | | Mixed | Down | | | | | | | |
| ***Lpar1*** | -0.22 | | 0.26 | -0.58 | | | -0.62 | | | Down | | | | | | | |
| ***Sox9*** | -0.17 | -0.13 | | -0.28 | | -0.05 | -0.24 | | | | -0.44 | | | 0.26 | 0.23 | | |
| ***Olfml1*** | -0.16 | | | -0.28 | | | -0.34 | | | | | -0.19 | | | | | |
| ***Atad2*** | -0.15 | -0.17 | | -0.20 | | | | | | | | -0.11 | | | | 0.06 | |
| ***Gdf11*** | -0.15 | | | | | | -0.12 | | | | | -0.23 | | | | | |
| ***Fads6*** | -0.13 | | -0.16 | | | | -0.15 | | | | | | | | | -0.07 | |
| ***S100b*** | -0.13 | -0.11 | 0.16 | | | | -0.32 | | | | | | | 0.14 | | | |
| ***Aqp4*** | -0.13 | | | | | | -0.73 | | Down | | -0.46 | | | 0.35 | 0.22 | 0.18 | 0.19 |
| ***Rbms1*** | -0.12 | | | -0.15 | | | -0.19 | Up | | | | 0.11 | -0.08 | | | | |
| ***Tek*** | -0.12 | | | -0.11 | | | -0.36 | | | | | | | | | -0.08 | |
| ***Mid1ip1*** | -0.11 | -0.06 | | -0.14 | | | -0.22 | | | | -0.18 | | | | | | |
| ***Elovl5*** | -0.11 | | | | | | -0.15 | | | | -0.31 | -0.65 | | 0.14 | 0.15 | | |
| ***Rnd2*** | -0.10 | -0.11 | | | | -0.07 | | | | | | | | -0.10 | | | |
| ***Anxa5*** | -0.10 | | 0.17 | | | | -0.14 | | | | | | -0.08 | | | | |
| ***Mtus1*** | -0.09 | | | | | | -0.22 | Mixed | Mixed | Down | | | | | | | |
| ***Slc25a15*** | -0.09 | -0.08 | | | | | | | | | | | | | | -0.11 | |
| ***Ankef1*** | -0.09 | -0.13 | | | | 0.25 | | | | | | | -0.05 | | | | |
| ***Kif1c*** | -0.08 | | | -0.13 | | | -0.08 | | | | -0.28 | | | | | | |
| ***Scaf8*** | 0.06 | 0.06 | | 0.09 | | | | | | | | | | 0.05 | | | |
| ***Phka2*** | 0.08 | | -0.18 | | | | | Up | | | | | -0.05 | 0.03 | | | |
| ***Myo1h*** | 0.12 | | | | | | 0.32 | | | | | | 0.05 | | | | |
| ***Chrd*** | 0.18 | | | 0.22 | | | 0.16 | | | | | 0.32 | | -0.06 | | -0.05 | |
| ***Xdh*** | 0.19 | 0.25 | | 0.25 | 0.33 | | 1.05 | | Up | | | | | 0.05 | | | |

***Table S3. Many of our DEGs may show similar effects to those observed in large studies characterizing the effects of stress-related disorders on the cortical transcriptome.*** *Provided in this table are the Log2FCs for DEGs that showed similar effects in the prefrontal cortex or cortex in either* ***1)*** *two or more human psychiatric datasets that resembled those observed following chronic stress, or* ***2)*** *two or more chronic stress datasets that resembled those observed in human psychiatric disorder. Throughout the table, blue highlighting indicates down-regulation in response to stress or glucocorticoids and pink highlighting indicates upregulation in response to stress or glucocorticoids. Italic text indicates nominal significance (p<0.05), non-italic text indicates significance in the original study (typically FDR<0.05). The studies referenced by the columns include 1) our current meta-analysis (Xiong et al.), 2) a meta-analysis of the effects of 10 days of CSDS on the PFC (Reshetnikov et al., 2022: n=56 subjects), 3) a smaller meta-analysis of the effects CSDS on the PFC in stress susceptible vs. stress resilient animals (Reshetnikov et al., 2022: n=18 subjects), 4) a smaller meta-analysis of the effects of 30 days of CSDS on the PFC (Reshetnikov et al., 2022: n=20 subjects), 5) a large meta-analysis of the effects of chronic stress more broadly defined (CSDS, CUMS, or chronic restraint) (Gururajan, 2022: n=101 subjects, only results surviving FDR<0.05 available), 6) a meta-analysis of the long-term effects of early life stress on the prefrontal transcriptome (Duan et al. 2025: n=89 subjects), 7) a large study of the effects of chronic daily corticosterone on the hippocampal transcriptome (Juszczak et al., 2025: n=48 subjects), 8-10) Comprehensive reviews of the literature characterizing the effects of chronic stress (medium duration or prolonged) and PTSD on the brain transcriptome (Stankiewicz et al., 2022), 11) Any of four meta-analyses of the effects of suicide in the frontal cortex (Piras et al. 2022), 12) The effects of Alcohol Abuse Disorder on the superior frontal cortical transcriptome (Gandal et al. 2018: n=32), 13-15) Meta-analyses of microarray data characterizing the effects of psychiatric disorders on the cortical transcriptome (Gandal et al., 2018) Gandal et al. 2018: Major Depressive Disorder (n=176), Bipolar Disorder (n=197), and Schizophrenia (n=314), 16-17) Meta-analyses of RNA-Seq data characterizing the effects of Bipolar Disorder (n=171) and Schizophrenia (n=384) in the frontal cortex (Gandal et al. 2018).*
